# Cellular heterogeneity of human fallopian tubes in normal and hydrosalpinx disease states identified by scRNA-seq

**DOI:** 10.1101/2021.09.16.460628

**Authors:** Nicole D Ulrich, Yu-chi Shen, Qianyi Ma, Kun Yang, D Ford Hannum, Andrea Jones, Jordan Machlin, John F Randolph, Yolanda R Smith, Samantha B Schon, Ariella Shikanov, Erica E. Marsh, Jun Z Li, Sue Hammoud

## Abstract

Fallopian tube (FT) homeostasis requires dynamic regulation of heterogeneous cell populations and is disrupted in infertility and ovarian cancer. Here we applied single-cell RNAseq to profile 53,376 FT cells from 3 healthy pre-menopausal subjects. The resulting cell atlas contains 12 major cell types representing epithelial, stromal and immune compartments. Re-clustering of epithelial cells identified 4 ciliated and 6 non-ciliated secretory epithelial subtypes, two of which represent potential progenitor pools: one leading to mature secretory cells, while the other contributing to either ciliated cells or one of the stromal cell types. To understand how FT cell numbers and states change in a disease state, we analyzed ~15,000 cells from a hydrosalpinx sample and observed shifts in epithelial and stromal populations, and cell type-specific changes in extracellular matrix and TGF-β signaling, underscoring fibrosis pathophysiology. This resource is expected to facilitate future studies to understand fallopian tube homeostasis in normal development and disease.

## Introduction

In mammals, the fallopian tube (FT) is a highly specialized narrow organ that is essential for the transport of gametes and embryos during unassisted fertilization. Anatomically, the tube can be partitioned into three segments with distinct functions: fimbriae, the finger-like structures responsible for oocyte capture after ovulation; ampulla, the site of oocyte fertilization by the sperm and the first 3 days of embryo development; and isthmus, which serves as the reservoir for sperm prior to fertilization, and through which the embryo is transported to the uterus for implantation (Barton et al. 2021; Crow et al. 1994).

Each segment of the fallopian tube contains both epithelial and stromal layers. The epithelial layer consists of four main types of cells: ciliated, secretory, peg and basal cells(Paik et al. 2015; Peters 1986). The ciliated cells (FOXJ1^+^/CAPS^+^), the most abundant and well defined, control gamete and embryo movement during fertilization and implantation (Paik et al. 2015; Barton et al. 2021). The secretory cells (PAX8^+^/KRT7^+^) produce and secrete factors required for fertilization and early embryo development. The peg and basal cells are less abundant. Although peg cells are often defined by EPCAM^+^/CD44^+^/ITGA6^+^, their biological functions are poorly understood (Paik et al. 2015; Peters 1986). *In vitro*, peg cells are capable of expanding and generating tube-like organoids consisting of both ciliated and secretory cells, suggesting that peg cells may have stem cell or progenitor properties (Paik et al. 2015). Basal cells are defined histologically by their round morphology with a cytoplasmic ring, and by expression of epithelial (EPCAM) and T-cell or resident T-cell markers, indicative of potential immune function (Peters 1986; Hu et al. 2020). Compared to the epithelial layer, the stromal layer contains a larger number of distinct cell types, including lymphocytes, macrophages, dendritic cells, mast cells, smooth muscle cells, fibroblasts, blood and lymphatic endothelial cells, and possibly Cajal-like cells (Ardighieri et al. 2014; Popescu, Ciontea, and Cretoiu 2007).

Although epithelial and stromal cell populations are present in all anatomic segments of the fallopian tube, their relative proportions may differ depending on hormonal states, menopause status, diseases such as ovarian cancer, and surgical disruption such as tubal ligation (Donnez 1985; Crow et al. 1994; Tiourin et al., n.d.; Paik et al. 2015; Ely and Mireille 2013; Gaitskell et al. 2016). For example, in fertile women, ciliated cells are most abundant in the fimbriae, and reach their highest numbers around the time of ovulation; but their numbers decrease during long-term progestin therapy (Donnez 1985). Conversely, ciliated cells are decreased in number in the fimbriae of post-menopausal women, but are restored to pre-menopausal levels with estrogen therapy (Crow et al. 1994). Similarly, after tubal ligation, the numbers of peg cells - presumed progenitors of the fallopian epithelium - decrease in the fimbriae. This decrease in peg cells was associated with a decreased risk of developing ovarian cancer in both pre- and post-menopausal women (Paik et al. 2015; Ely and Mireille 2013; Gaitskell et al. 2016; Tiourin et al., n.d.), which led to the hypothesis that peg cells may be the cells of origin for high-grade serous ovarian cancer (Hu et al. 2020; Dinh et al. 2021).

Additionally, changes in tubal epithelium composition can also lead to a form of female infertility known as tubal factor infertility, which in some women can be associated with hydrosalpinx - an enlarged and fluid-filled fallopian tube (Sapmaz et al. 2019; Ajonuma et al. 2005; Yohannes et al. 2019). For these women, tubal factor infertility treatment involves the surgical removal of hydrosalpinx, followed by *in vitro* fertilization and embryo transfer (Strandell 2000; Johnson et al. 2010; Sapmaz et al. 2019). We currently have a limited understanding of the etiology of hydrosalpinx, and we lack alternative therapies (other than surgical removal) that could retain tubal function and allow unassisted fertility. Hence, gaining a greater understanding of the cellular heterogeneity of the healthy fallopian tube and the key changes in diseases can help us develop better therapeutic strategies for treating reproductive organ pathologies.

In this study, we applied single-cell RNA-sequencing (scRNA-seq) to analyze 53,376 unselected cells from 9 specimen of human fallopian tubes from 3 healthy subjects. This unbiased characterization of >53K single cells allowed us to define 12 major cell types in healthy FT. Through iterative re-clustering of the two epithelial cell types, we identified 4 ciliated epithelial (CE) and 6 non-ciliated secretory epithelial (NCSE) subtypes. We validated their presence in native tissues using immunofluorescence of marker proteins in FFPE sections. Importantly, we identified two of these subtypes as likely progenitor cells that could play a key role in fallopian tube homeostasis. To better understand how cell populations change in hydrosalpinx, we sequenced ~15,000 cells in a surgically removed FT sample from a patient with this condition. In hydrosalpinx FT, several cell populations, including immune cell and stem cell/progenitor populations, differed significantly from the healthy state. In addition, extracellular matrix (ECM) genes were altered, and TGF-β signaling pathways were upregulated, indicative of tissue fibrosis. Overall, the catalog of cell types, subtypes, and specific markers represent a new cell atlas of healthy human fallopian tube and identify potential progenitor cells responsible for the highly regulated maintenance of this dynamic organ. The comparison with samples from a diseased FT provides the first glimpse of the pathophysiology of hydrosalpinx, for both altered cell number and cell type-specific dysregulation of ECM and TGF-β genes. These insights may help guide the development of new therapies as alternatives to surgical intervention.

## RESULTS

### Systematic analysis of fallopian tube from 3 healthy individuals revealed 12 major cell types

To ensure that our tissue dissociation protocol was sufficiently robust to include the full range of FT cell heterogeneity, we dissected a benign surgical sample (FT1) into three segments, fimbria, isthmus and ampulla. We collected live cells from both the epithelial and the stromal compartments and performed scRNA-seq analysis (**Figure 1A**). Through unsupervised clustering of 3,722 cells from isthmus, 4,944 cells from ampulla, and 1,861 cells from fimbria we identified an initial set of 11 molecular clusters from each segment, which were highly concordant in their distributions across the 3 segments (**Supplemental Figure 1A**) and in their expression profiles (**Supplemental Figure 1B**). By analyzing cluster-specific RNA markers (**Supplemental Table 1**) we annotated the 11 clusters and found that they represent all known major cell types in FT, thus validating that our dissociation protocol captured the expected cellular diversity of the organ. Next, we extended the same analysis to 3 tubal segments from a second benign surgical sample (FT4), and from a cadaveric sample with normal anatomy (FT2) (see their clinical features in **Supplemental Table 2A**). Systematic comparisons of the clusters found independently in FT1, FT2 and FT4 confirmed high individual-to-individual correlations of matched cell types (**Supplemental Figure 1C**) and tissue composition (**Supplemental Figure 1D, E**), indicating that the heterogeneous cell populations have been reproducibly observed in three independent experiments, which included both surgical (FT1, FT4) and cadaveric (FT2) samples.

**Figure 1.**
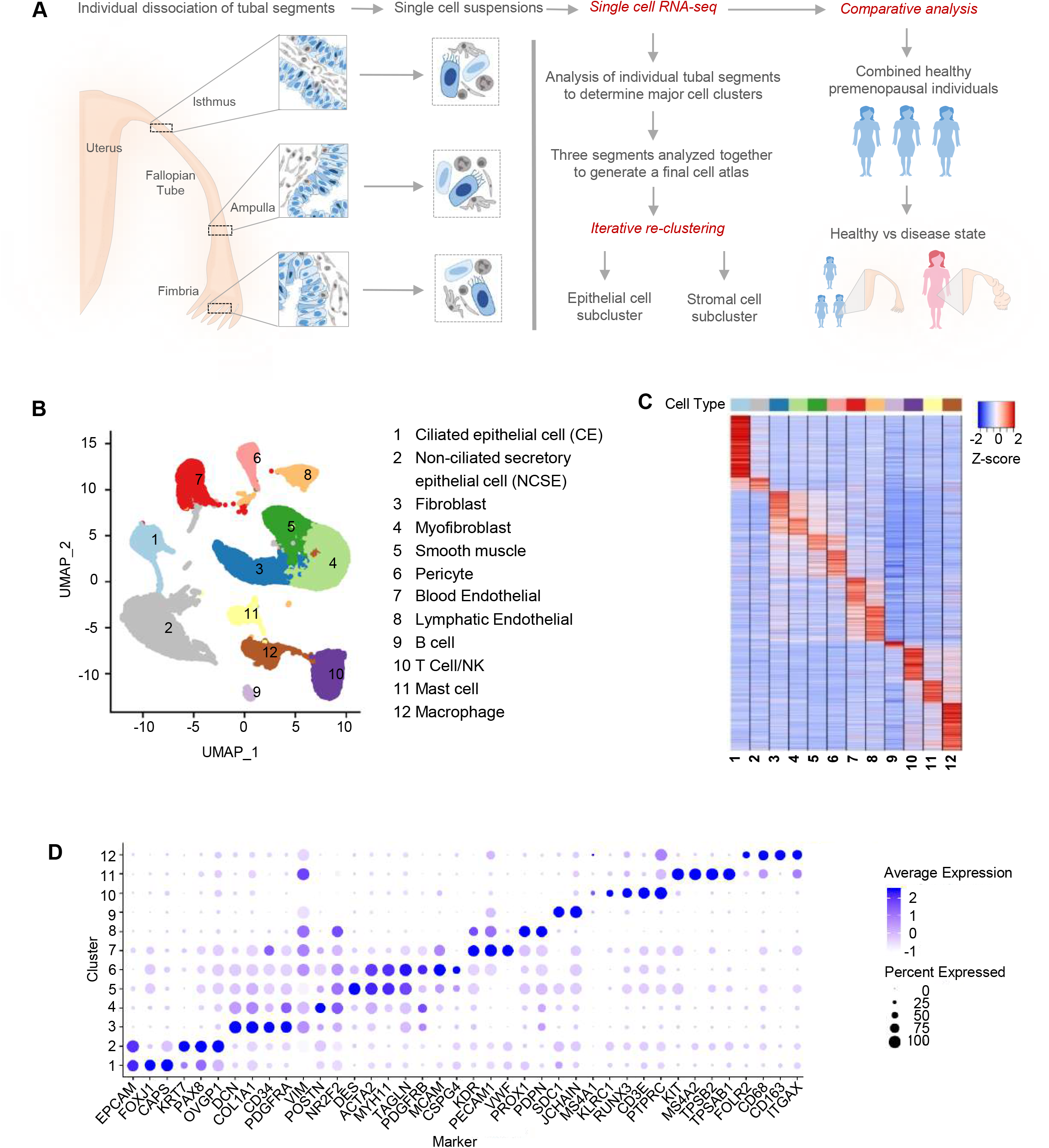
Major cell types and markers identified from single-cell RNA-Seq analysis of healthy human fallopian tubes. A. Overview of the study, including the fallopian tube sections from which the samples were taken, and the data collection and analysis processes. B. Identification of 12 major cell types from global clustering of 53,178 high-quality cells from three healthy subjects, visualized in UMAP space. C. Marker gene expression pattern in the 12 major cell types, with values for each gene averaged within each cell type (i.e., the centroid), then standardized over the 12 centroids. D. Average expression level and prevalence of selected major markers used to annotate the 12 major fallopian tube cell types. Details are in Table S3.

To exploit the full power of the entire data we pooled the total set of 53,376 pass-QC cells from the 3 subjects, at an average of ~2.6K detected genes per cell. Through batch-correction and re-clustering (see **Computational Methods**) we defined a stable set of 12 clusters (**Figure 1B**), annotated them by marker genes (**Figure 1C–D**), with their average expression levels provided in **Supplemental Table 3**. The three individuals contributed comparably to each cell type (**Supplemental Figure 1D, E**). Mainly by relying on known marker genes (**Figure 1D**) we identified two major epithelial cell types (EPCAM), consisting of ciliated cells (FOXJ1, CAPS) and non-ciliated cells (PAX8, KRT7, OVGP1). Within the stromal compartment we assigned 6 cell types: fibroblast (DCN, COL1A1, PDGFRA), myofibroblast (POSTN, PDGFRB, ACTA2), smooth muscle (DES, MYH11, ACTA2), pericyte (MCAM, CSPG4), blood endothelial (VWF, KDR, PECAM1), and lymphatic endothelial (PDPN, SCD1). Lastly, we identified 4 immune cell types: B cell (JCHAIN, SDC1), T/NK cell (RUNX3, CD3E, KLRC1), mast cell (KIT, MS4A2, TPSB2), and macrophage (CD68, ITGAX, CD163). The only cell type previously reported in the fallopian tube (Popescu, Ciontea, and Cretoiu 2007) that we failed to identify is the Cajal-like cell. This may be due to their long cytoplasmic processes and small cell body, causing these interstitial cells to be damaged or lost during dissociation. Despite this, our collection of ~53K high-quality single cells portrays the functional and molecular diversity of the premenopausal human fallopian tube and covers all known major cell types of the organ. In the following, we describe focused clustering of the ciliated cells, and separately the non-ciliated epithelial cells, to uncover finer subtypes and their potential lineage relationships.

### Identification and characterization of four ciliated epithelial subtypes

By re-clustering the ~2.5K ciliated epithelial cells identified in the global analysis (**Figure 1B**) we identified four molecularly defined subtypes (**Figure 2A**), which were positive for FOXJ1 and CAPS (**Figure 2B**) and were contributed comparably by the three individuals (**Supplemental Figure 2A**). We designated them as CE1-1 through CE1-4. Based on subtype-specific RNA markers, CE1-1 cells are likely mature, multi-ciliated cells, as they express genes encoding proteins associated with motile cilia and flagella (CFAP157, CFAP73, CFAP52, and CFAP126), dyneins (DYNLL1, DNAI1, DNAI2, DNAH11, KIF21A), radial spike proteins (RSPH1, 4a, and 9), intraflagellar transport proteins (IFT57), and calcium sensing and binding proteins (CAPS, CALM1) (examples in **Figure 2C**; full list in **Supplemental Table 4**). CE1-2 cells express a relatively high level of ribosomal protein-related genes (RPL12, RPL21, RPL31, PRS18 and RPL26) and those involved in apoptosis (**Supplemental Table 4**). Despite the abundance of ribosomal proteins, these cells have a notably lower number of detected transcripts than other CE subtypes (**Supplemental Figure 2B**), prompting us to ask whether these cells are damaged or otherwise of lower quality. The relative abundance of mitochondria-encoded transcripts, which tends to be higher in damaged cells, was the lowest in CE1-2 (**Supplemental Figure 2C**).Furthermore, we estimated the contribution of ambient RNA to each cell by defining an *in-silico* transcriptome profile for damaged cells – the “soup”– using the thousands of cells that fell below the minimal number of detected genes, and then calculating the “distance” of every cell to the centroids of (1) the 4 CE subtypes and (2) the soup. CE1-2 cells are distinct from “soup” (**Supplemental Figure 2D**), thus confirming that CE1-2 cells are naturally existing and undamaged. Given their low RNA content and gene markers for apoptosis and p53 pathway (**Supplemental Table 4**), we speculate that CE1-2 cells are either senescent or executing programmed cell death. The RNA levels of senescence markers such as CDKN2A and CDKN2B did not vary across the four ciliated clusters (not shown); however, they may still be regulated at the protein level. Indeed, images from the human protein atlas database suggest that a rare number of CDKN2A cells are scattered in the human fallopian tube epithelium (**Supplemental Figure 2E**).

**Figure 2:**
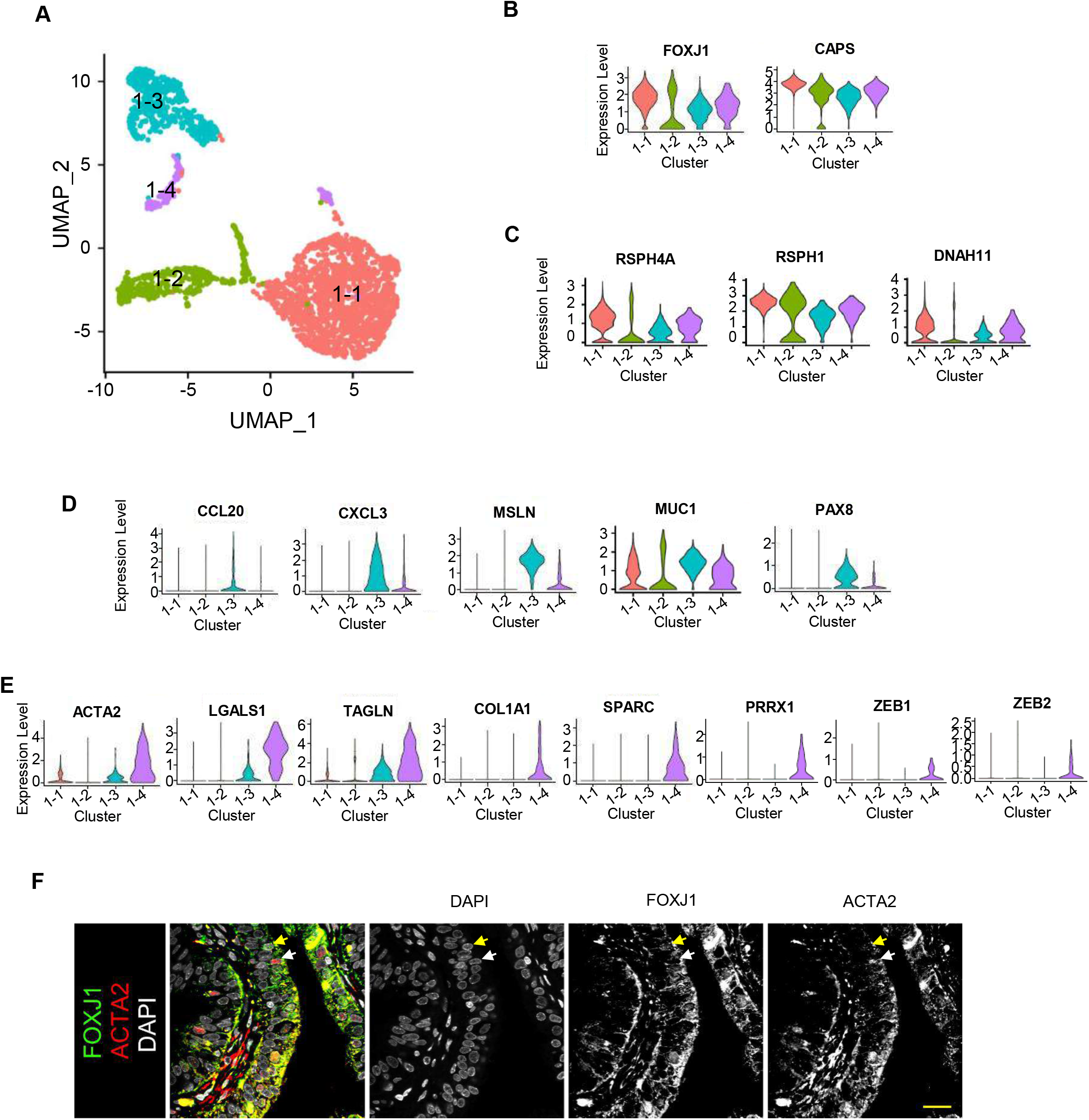
Subtypes of ciliated cells. A. Focused re-clustering 2,510 ciliated cells of healthy fallopian tubes identifies 4 subtypes, 1-1 to 1-4, shown in UMAP space. B-E. Expression levels of selected markers used to identify the 4 ciliated cell subtypes, with general markers for ciliated cells are shown in B; markers for 1-1 and 1-2 in C; those for 1-3 in D; and those for 1-4 in E. F. Immunofluorescence (IF) staining of the fallopian tube epithelium using antibodies against unique markers for ciliated cell subtypes. Yellow arrow: FOXJ1^+^ cell. White arrow: ACTA2^+^ cell.

CE1-3 cells express genes involved in antigen processing (CD74), presentation (HLA-DMA, HLA-DRB1), and chemokine/cytokines (CCL20, CXCL1-3) (**Supplemental Table 4;** examples in **Figure 2D**). CE1-4 cells express extracellular matrix genes COL1A1, TAGLN, LGALS (**Figure 2E**), and markers for epithelial-mesenchymal transition (EMT), which were also observed in NCSE2-2 (see below). We validated the presence of some CE subtypes *in vivo* with immunofluorescence data (**Figure 2F**). The FOXJ1/ACTA cells likely correspond to mature CE1-1 and CE1-3 cells, while the FOXJ1/ACTA2 cells likely represent CE1-4, a precursor to mature ciliated cells.

### Focused analysis of non-ciliated epithelial cells identified six subtypes

Re-clustering of the ~14,000 non-ciliated epithelial cells from FT1, FT2 and FT4 identified 6 molecular subtypes (**Figure 3A**), contributed comparably by the three subjects (**Supplemental Figure 3A)**, and are named NCSE2-1 through NCSE2-6. All six subtypes express classic secretory cell markers EPCAM, PAX8, and KRT7 (**Figure 3D**). Using differentially expressed markers (**Supplemental Table 5; Figure 3B**) we deduced their potential functional roles.

**Figure 3:**
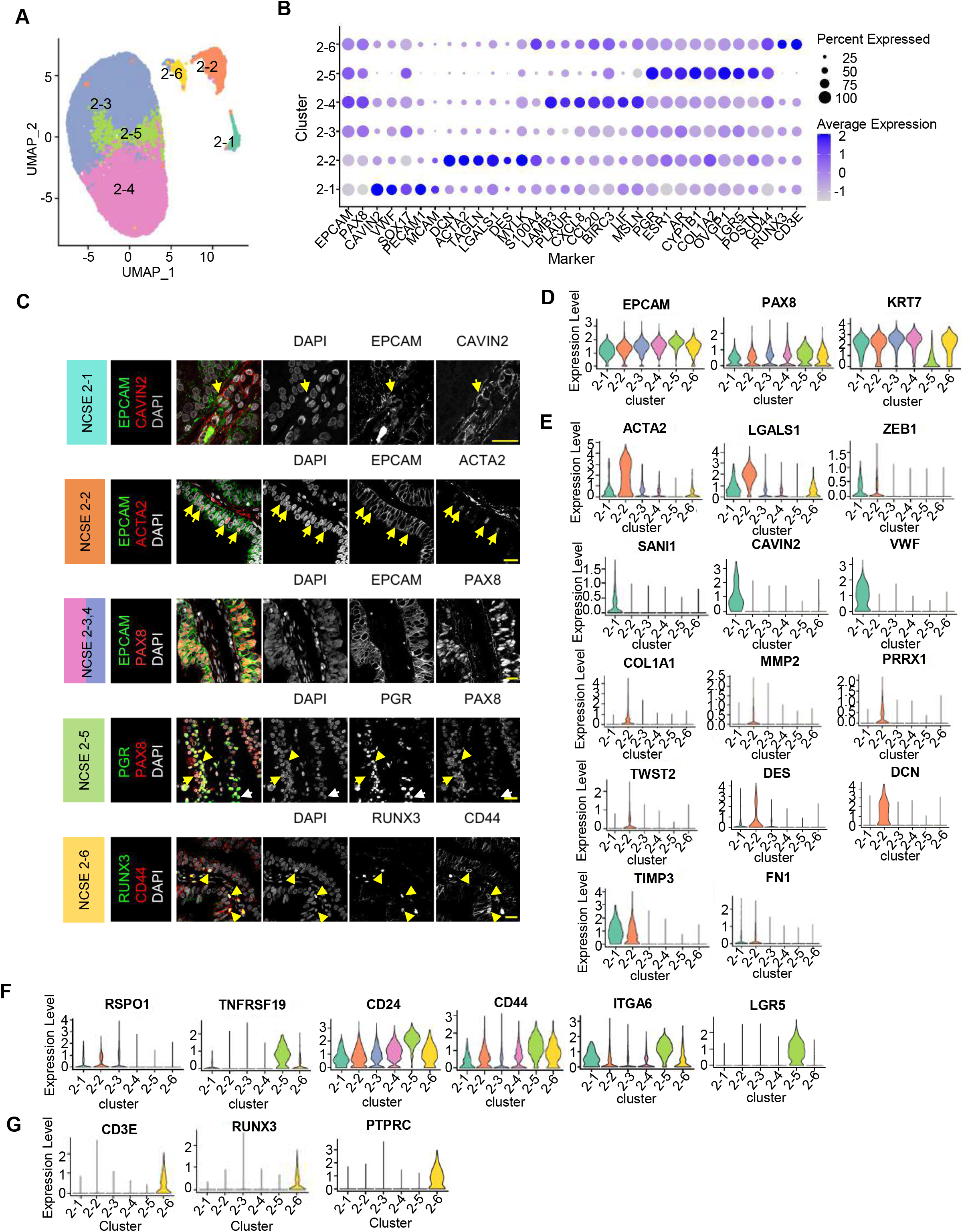
Subtypes of non-ciliated secretory epithelial cells. A. Focused re-clustering 14,002 non-ciliated secretory epithelial cells of healthy fallopian tubes identifies 6 subclusters, 2-1 to 2-6, shown in UMAP space. B. Average expression levels and prevalence of major markers used to identify the 6 non-ciliated secretory epithelial cell subtypes. C. IF staining using antibodies against unique markers for non-ciliated secretory epithelial cell subtypes. D. Expression levels of common markers for non-ciliated epithelial cells. E-G. Expression levels of selected markers used to identify the 6 subtypes. E. Markers for NCSE 2-1 and 2-2. F. Makers for NCSE 2-3 (RSPO1) and NCSE 2-5. H. Markers for NCSE 2-6.

NCSE2-1cells account for 1.6-2% of the NCSEs (**Supplemental Figure 3B**), and express EMT markers ACTA2, ZEB1, SNAI, LGALS1, SPARC, and endothelial cell markers CAVIN2, PECAM, VWF (**Figure 3B, E**). To examine whether the epithelial, EMT, and endothelial cell markers are co-expressed in a single cell, or in different cells, we evaluated pairwise co-expression patterns between EPCAM, ACTA2, and CAVIN2, over the individual NCSE cells (**Supplemental Figure 3C**) and confirm that EPCAM-ACTA2 and EPCAM-CAVIN2 are co-expressed in NCSE2-1 cells. Consistent with the rarity of NCSE2-1 in scRNA-seq data, EPCAM and CAVIN2 proteins are colocalized in a small fraction of cells *in vivo* (**Figure 3C**, first row). These EPCAM/CAVIN2 cells do not appear to be intercalated within the epithelium, rather they abut the epithelium and are close to blood vessels, suggesting that these cells may be an epithelial cell population in the process of transitioning to endothelial cells.

Like NCSE2-1, NCSE2-2 cells express EMT markers ACTA2, COL1A1, MMP2, PRRX1, TWIST2, DES, DCN, FN1, ZEB1, TIMP3, LGALS1, SPARC (**Figure 3C, E; Supplemental Table 5**), but lack any terminal differentiation markers of epithelial cells, consistent with a progenitor-like cell. NCSE2-2 cells account for 5-8% of NCSE cells (**Supplemental Figure 3B)**. To validate their presence *in vivo*, we immunostained all three segments of the fallopian tube for EPCAM and ACTA2 and found that EPCAM/ACTA2 double-positive cells are relatively abundant and intercalated in the epithelium as clusters (**Figure 3C**, second row). The identification of this population with EMT marker expression led us to examine their distribution in a larger collection of 11 benign surgical samples varying by age and hormonal status (see their demographic and clinical information in **Supplemental Table 2B)**. By immunofluorescence-based counting of EPCAM/ACTA2 double positive cells in pre- and post-menopausal samples, we found that these cells were present in all 3 anatomic segments, although their frequency varied across subjects representing different phases of the cycle and varied by anatomic segments (**Supplemental Figure 3D**). Despite the variability, samples from post-menopausal women appear to have fewer EPCAM/ACTA2 cells, except for one woman known to be using vaginal estrogen therapy (boxed). To assess if these cells might be directly regulated by estrogen or progesterone, we immuno-stained the tissue sections for ACTA2 and estrogen receptor (ESR) or progesterone receptor (PGR) and found very few ACTA2 cells co-expressing ESR or PGR (**Supplemental Figure 3E**), consistent with the relatively low ESR and PGR RNA expression levels in NCSE2-2 (**Supplemental Figure 3F**). While these results suggest that the EPCA/ACTA cell population is not directly regulated by estrogen or progesterone, it remains possible that the frequency of these cells is indirectly modulated by estrogen or other hormones, such as androgen.

NCSE2-3 to 2-5 are the most abundant subtypes, collectively comprising ~80% of NCSE (**Supplemental Figure 3B**). In addition to expressing PAX8, KRT7 and OVGP1, NCSE2-3 cells can be distinguished by expression of SOX17 and WNT-signaling regulator RSPO1 (**Figure 3F**). NCSE2-4 cells express genes related to ovarian cancer (LAMB3, MSLN), and LIF (**Figure 3B**), an interleukin 6 class cytokine essential for normal embryo implantation (Hassan et al. 2006; Ribeiro et al. 2018). Notably, NCSE2-5 cells express stem/progenitor markers such as LGR5, CD24, and TNFRSF19 (**Figure 3B, G**) (Ng et al. 2014; La Manno et al. 2018; Lamouille, Xu, Jian, and Derynck, Rik 2014; Mabbott et al. 2013; Blanpain, Horsley, and Fuchs 2007), as well as CD44 and ITGA6 (**Figure 3F**), markers previously used to define the peg cell population. This population may be hormonally regulated, due to its high expression levels of hormone receptors ESR, PGR, and AR (**Supplemental Figure 3G**), and CYP1B1 (**Figure 3B**), a P450 enzyme known to metabolize 17β-estradiol to a 2- and 4-OH estradiol.

Finally, NCSE2-6 cells express lymphoid lineage markers RUNX3, CD3E, PTPRC (**Figure 3B, H**), consistent with a basal or resident memory T-cells identity. The EPCAM^+^/RUNX^+^/CD44^+^ cells are round, and basally located with a cytoplasmic halo (**Figure 3C**), likely corresponding to the previously described basal cells (Peters 1986). In contrast to NCSE2-2, the NCSE2-6 cells appear with consistent frequencies across premenopausal individuals in different cycle phases (**Supplemental Figure 3D**), and only varied in post-menopausal women.

Taken together, our analysis of ~16,500 NCSE and CE cells provided a fine-grained catalog of cellular diversity of the fallopian tube epithelium. The large number of high-quality cells allowed us to delineate 10 subtypes that would have been impossible to separate using previous methodology.

### Subtle transcriptomic differences are observed across anatomic segments

Although cell type composition is largely similar across the fallopian tube segments, we explored if there are between-segment transcriptomic differences within each cell type. We performed differential expression analysis for each defined cell type, and generally found very few genes with significant difference between segments (**Supplemental Table 6**). Nonetheless, the differentially expressed genes that did emerge reflect subtle differences between segments. For ciliated cells, the genes that show a higher expression in isthmus include those encoding the progesterone receptor (PGR), blood clotting component Fibrinogen Alpha Chain (FGA), apoptosis regulator Immediate Early Response 3 (IER3), receptor tyrosine kinase regulator Leucine-rich alpha-2-glycoprotein 1 (LRG1), and metabolic regulator thioredoxin-interacting protein (TXNIP) (examples shown in **Supplemental Figure 5A**). The higher expression of PGR in isthmus is notable, as progesterone is known to be required for the highly coordinated movements of cilia in the isthmus, the site of fertilization and embryo transfer to the uterus for implantation (Barton et al. 2021). For both CE and NCSE populations, the expression of stemness markers ALDH1A1 and ALDH1A2 tends to be higher in fimbriae. In NCSEs, some ovarian cancer-associated markers and genes regulating cellular redox homeostasis (e.g., BIRC3, FGA, IER3) are elevated in fimbriae (**Supplemental Figure 5B; Supplemental Table 6**), supporting the hypothesis that the cell of origin of ovarian cancer resides in the fimbriae, and highlighting the increased oxidative stress in the distal tube in response to ovulation (Sowamber et al. 2020). Further, the non-ciliated cells express higher levels of inflammatory cytokines CXCL1 (IL-1), CXCL8 (IL-8), and the sperm chemoattractant CCL20 in the fimbriae (**Supplemental Figure 5A**), consistent with previous reports (Sowamber et al. 2020). Taken together, although transcriptomic differences are subtle, the genes identified point to functional differences across segment compartments, servings as a baseline for future studies exploring the genes dynamically regulated in response to disease or during pregnancy.

### Identification of six stromal subtypes in human fallopian tube

The global clustering (**Figure 1B**) involved ~26,400 stromal cells and identified six stromal cell types (re-projected in **Figure 4A**), which are evenly contributed by the three subjects (**Supplemental Figure 4A**). Cluster-specific genes (**Figure 4B**) allowed annotation of fibroblast (DCN, PDGFRA), myofibroblast (PRRX1, POSTN), and smooth muscle cell (ACTA2, MYH11); and we validated their presence *in vivo* using immunofluorescence or immunohistochemistry of representative markers (**Supplemental Figure 4B**). These three cell types appear to be the most similar to each other (**Figure 4A**) and have the highest expression of hormone receptors among the stromal cells (**Figure 4C**), consistent with their function being influenced by hormonal status. The next three cell types are pericyte (MCAM, PDGFRB, CSPG4), blood endothelial (VWF, SELE), and lymphatic endothelial (PPROX1, LYVE1, CAVIN2) cells (**Figure 4B),** which we identified *in vivo* using MCAM, VWF, and CAVIN2 immunocytochemistry **(Supplemental Figure 4B**).

**Figure 4:**
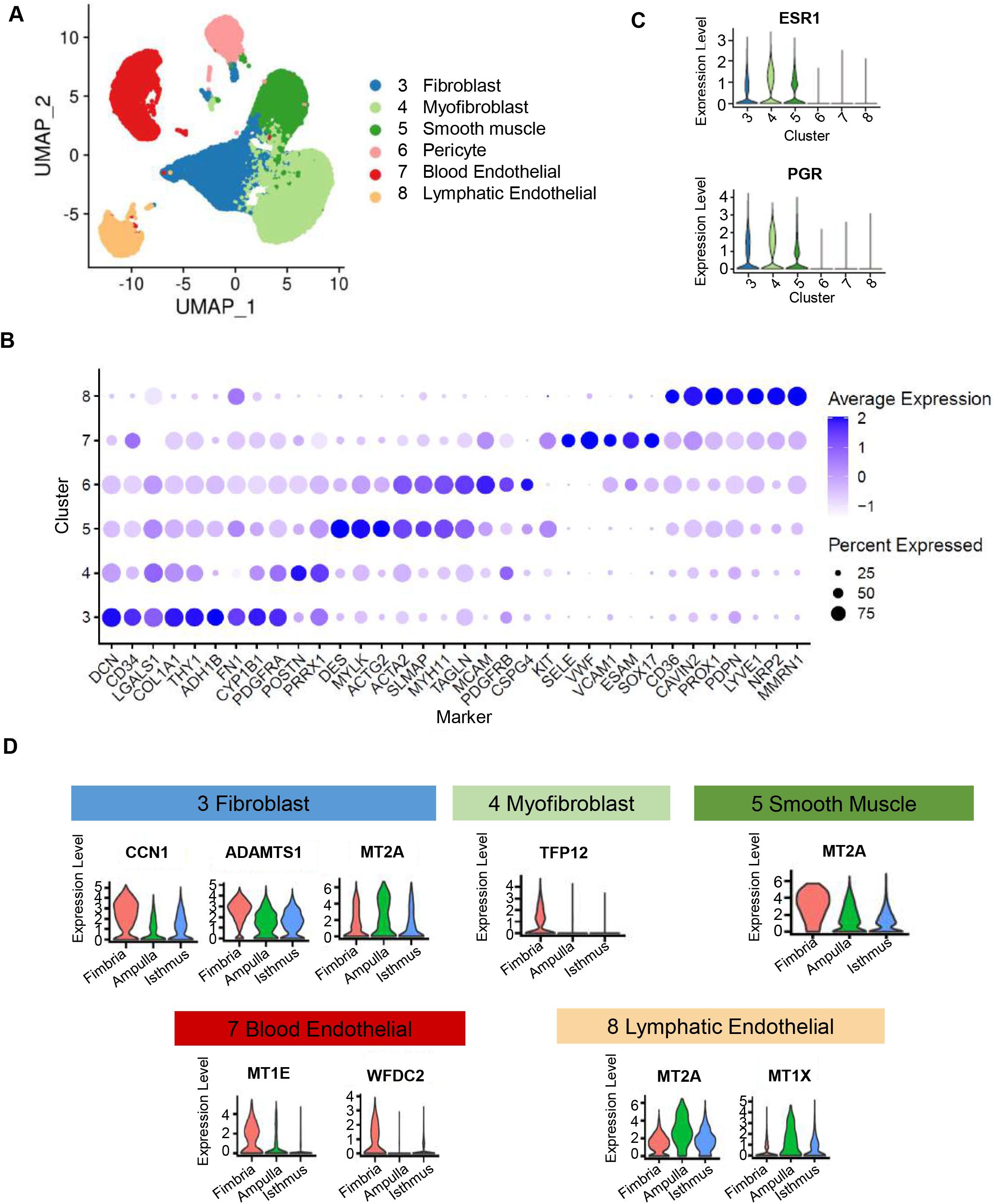
Stromal cell classification. A. Visualization of 42,911 stromal cells in a UMAP projection of only the stromal cells, colored by the 6 stromal clusters from global clustering shown in Fig.1B B. Average expression levels and prevalence of major markers used to annotate the stromal cell types. C. Expression levels of genes encoding hormone receptors, ESR1 and PGR, in the 6 stromal cell types. D. Expression levels of select markers differentially expressed across the three fallopian tube segments for different stromal clusters.

Like the epithelial cells, the stromal cell types display subtle differences in gene expression across segments (**Supplemental Table 6)**. For fibroblast cells, CCN1, a matrix-associated angiogenesis and cell proliferation regulator, is higher in the fimbriae, as expected given their dense vascularity. Additional genes that show segmental differences in specific cell types include serine protease inhibitor TFPI2 (myofibroblasts), metalloproteinase ADAMTS1 (fibroblast), stress proteins metallothionein 2A (MT2A) (smooth muscle), metallothionein 1X MT1X (lymphatic endothelial), and pro-angiogenic and immunosuppressive STAT3 pathway regulator WFDC2 (HE4) (expressed in pericytes and blood endothelial). Many of these genes are involved in ECM deposition or remodeling, suggesting possible differences in ECM composition across the tube. Taken together, the gene expression variation of the stromal cell populations across segments reveals a deeper complexity to gene regulation than previously appreciated.

### Identification of multiple progenitor populations that possibly contribute to epithelial and stromal cell homeostasis

Marker genes of NCSE2-5 and NCSE2-2 suggest that these populations may be progenitor cell populations in the fallopian tube epithelium. Next, we asked whether these populations are (1) part of a single, continuous developmental trajectory, (2) can contribute to different epithelial cell subtypes, or (3) can contribute to some of the stromal cell populations. As shown above, cluster analysis (**Figure3A**) revealed a discontinuous distribution, such that NCSE2-2 is not directly connected to other NCSE subtypes, and NCSE2-5 is linked to 2-3, 2-4, but not with other populations. To overcome the limitations of cluster analysis, we carried out RNA “velocity” analysis, which took advantage of the unspliced transcripts in scRNA-seq data to compare with the cell’s spliced transcripts (La Manno et al. 2018). The ratio of unspliced to spliced transcripts for each gene represent the difference between nascent and mature RNA and is used as a proxy measurement for the cell’s “velocity” in its differentiation trajectory, as depicted by the vectors in UMAP plots. First, when we jointly analyzed the six NCSE subtypes and the CE cells (**Figure 5A**), the velocity vectors for NCSE2-5 cells point unidirectionally towards NCSE2-4, a mature secretory cell population, while making no contribution to other epithelial cell types, suggesting that the LGR5^+^ NCSE2-5 cells are likely precursors for mature secretory cells. In contrast, NCSE2-2 cells, those with quasi-EMT features, contribute to NCSE2-1 cells, an intermediate between epithelial cells and endothelial cells (**Figure 5B**). This intermediate population subsequently projects to the blood endothelial cell population (cluster 7) in the stroma – raising the possibility that an epithelial progenitor undergoes complete EMT to give rise to a stromal cell type (**Figure 5B**).

**Figure 5.**
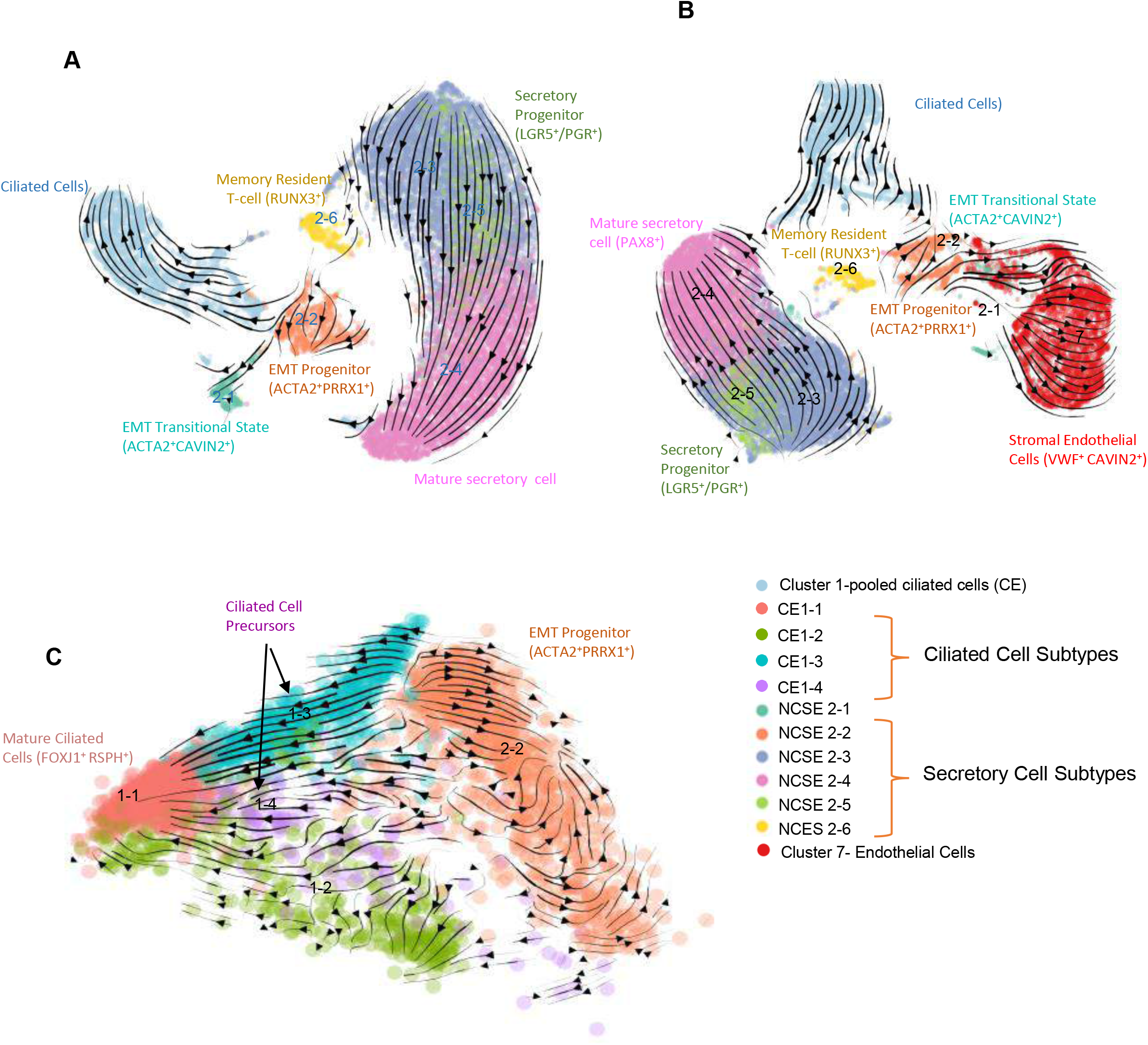
Velocity analysis that uncovers two potential progenitor populations. A. Velocity for CE and 6 subtypes of NCSE in UMAP view. B. Velocity for 4 subtypes of CE, NCSE 2-2 and Blood endothelial cells (7) in UMAP view. C. Velocity for 4 subtypes of CE and NCSE 2-2 in PCA view.

In addition to contributing to the stroma, NCSE2-2 cells also feed into the ciliated cells; thus, they are potentially responsible for the replenishment of ciliated cells. We repeated the velocity analysis by combining the NCSE2-2 cells with the ciliated cells, while distinguishing the 4 CE subtypes (**Figure 5C**). NCSE2-2 cells project to CE1-4, which in turn project to CE1-1, suggesting that CE1-4 are descendants of NCSE2-2, while at the same time represent precursors of CE1-1, the more mature ciliated cells. Marker genes of CE1-4 also supports its role as an intermediate population: they retain residual expression of NCSE lineage marker PAX8 (**Figure 2D**), express lower levels of EMT markers and lower levels of terminally differentiated ciliated cell markers such as FOXJ1, CAPS, RSPH genes, DNAH11 (**Figure 2B–E**). In addition, we also observe strong flow from CE1-3 to CE1-4, suggesting a parallel route to CE1-4 cells, although the source of CE1-3 cells remains unclear.

In sum, our trajectory analysis suggests that the fallopian tube has multiple progenitor populations that maintain multiple epithelial and stromal cell subpopulations in this dynamically regulated organ. Future spatial mapping and lineage tracing studies are needed to fully validate these putative developmental relationships.

### The hydrosalpinx tube had altered cell type proportions and cell states

In ultrastructure studies, the hydrosalpinx tube appears as a denuded epithelium with loss of cilia and microvilli, containing disordered edematous stroma with atrophic and distorted muscle fibers, and engorged blood vessels (Ajonuma et al. 2005). To investigate the cellular changes accompanying the abnormal structure and function of hydrosalpinx we performed scRNA-seq on a surgically removed hydrosalpinx tube from an individual (FT3) with tubal factor infertility and without evidence of endometriosis. Global clustering of the 15,571 cells from the sample reveals the same 12 clusters found in the healthy tubes, with no additional cell types (**Figure 6A**). Comparison of the relative cell number across the 12 cell types found that the hydrosalpinx tube had fewer ciliated and non-ciliated epithelial cells than the healthy tubes (**Figure 6B**). Among the stroma and immune cell types, fibroblast and macrophage populations are expanded, while the smooth muscle, pericyte, and endothelial cell populations are reduced (**Figure 6B**), indicating drastic tissue remodeling in the disease state.

**Figure 6:**
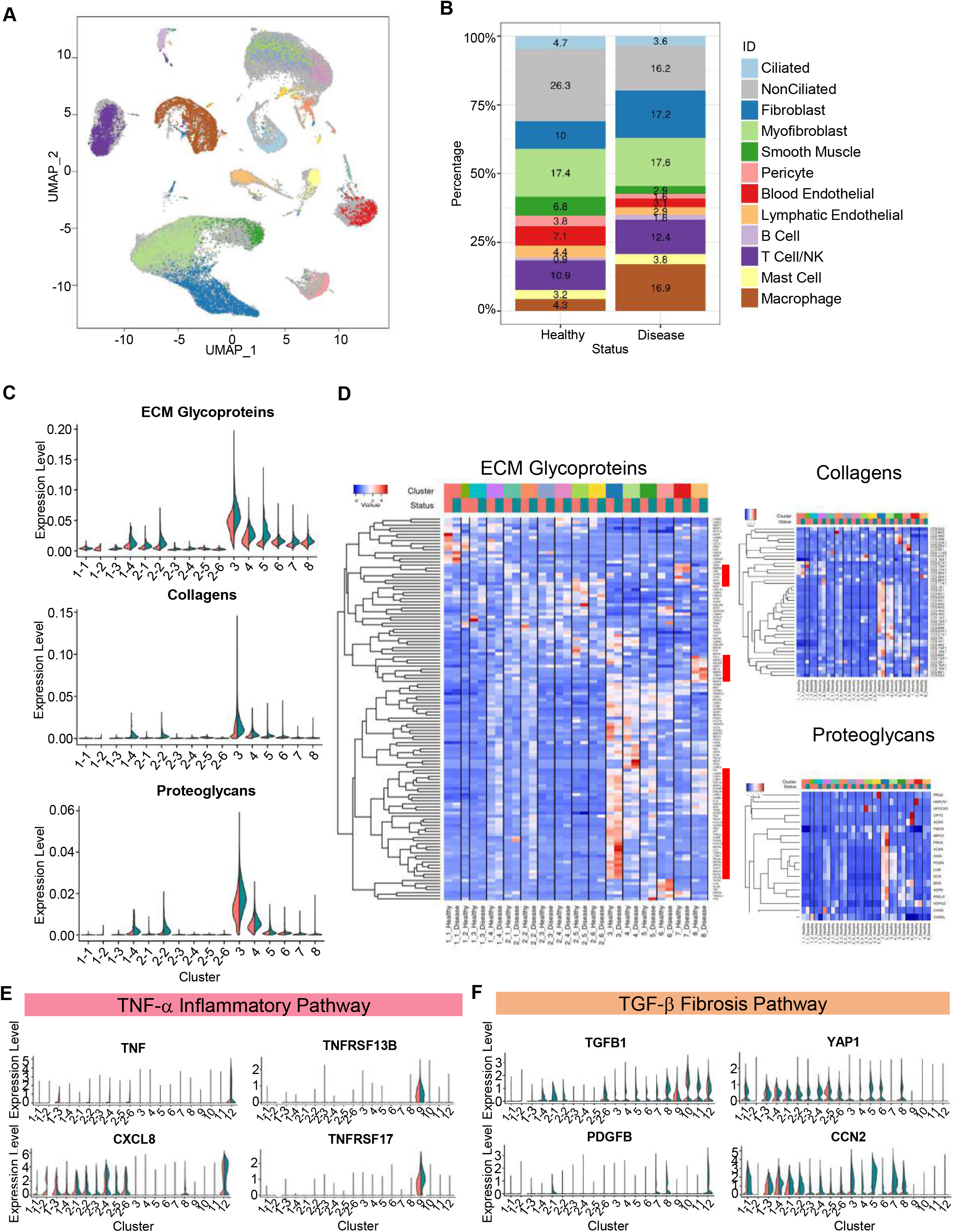
Comparison of epithelial and stromal populations between healthy and disease status. A. Visualization of cells from a diseased fallopian tube sample (FT3) in a joint UMAP projection with cells from the 3 healthy fallopian tube samples (FT1, FT2 and FT4), colored by supervised assignment of FT3 cells into clusters found for the healthy samples (colored as grey background). B. Composition of the 12 cell types, compared between the 3 healthy samples and the disease sample. C. Expression levels of three ECM-related gene sets, for the 4 CE subtypes. 6 NCSE subtypes, and 6 stromal cell types, compared between healthy (red) and disease (blue) samples. D. Heatmaps of expression levels of three sets of genes, for ECM glycoproteins, collagens, and proteoglycans, respectively, compared across cell types and between healthy (red) and disease (blue) samples. Average expression of each gene is calculated for each cell type (i.e., the centroid), then standardized over the centroids shown. E, F. Expression levels of genes in TNFα pathway (E) and TGFβ pathway (F), compared across cell types.

Next, we performed differential expression analysis between the healthy and disease samples within each epithelial and stromal cell type. Many of the genes upregulated in the disease state were involved in cell growth and proliferation (see **Supplemental Table 7**); and pathway analyses highlighted ERK1/ERK2 and the WNT signaling pathways (**Supplemental Table 8**). Consistent with the gene and pathway results, the percentage of MKI67^+^ cells were higher in the diseased than in the healthy tissue (**Supplemental Figure 6A**), reflecting a compensatory mechanism of higher proliferation in multiple epithelial subtypes after the loss of CE and NCSE cells (**Figure 6B**). Among the differentially expressed genes, many markers previously associated with adult somatic and cancer stem cell populations (ALDH1 and 2, THY1, LGR5, PDGFRA, PDGFRB), and EMT markers (ZEB1, LIF, POSTN, LGALS, MMP2, ACTA2) are elevated in the disease state (**Supplemental Figure 6B**). In particular, the expression of LGR5, which marks NCSE2-5 – an epithelial cell progenitor, is lower in the diseased sample, whereas levels of THY1 and PDGFRA, markers of mesenchymal stemness, and ALDH, are higher in the diseased state (**Figure 3B, Supplemental Figure 6B**). Importantly, these markers are generally higher in the NCSE2-2 progenitor population, suggesting that the balance between the two progenitor populations, NCSE2-5 and NCSE 2-2, is altered in the hydrosalpinx tube. Hydrosalpinx by itself has not been recognized as an ovarian cancer risk factor; but our data revealed changes in EMT and progenitor signature, thus possibly consistent with an elevated risk of cell transformation and tumorigenesis. This is supported by an increased expression of ovarian cancer markers in the fallopian tube (**Supplemental Figure 6C**).

In addition to the differences in progenitor markers, the differentially expressed genes revealed changes in the complement pathway, cellular redox, and WNT signaling pathways (**Supplemental Table 8**), consistent with previous analyses of hydrosalpinx. Specifically, the development of tubal disease and ultimately hydrosalpinx are associated with an inflammatory response to urogenital pathogens via the TNF-α pathway (Xu et al. 2019; Rasmussen et al. 1997; Morales et al. 2006). Proteomics analysis of tubal fluid uncovered abnormalities in the complement and redox pathways (Yohannes et al. 2019). We did not observe in the disease state an increase in TNF,TNFSF13B, or its receptor TNFRSF17 (**Figure 6E**), reported to be elevated in cases of salpingitis, or in active infections of the tube (Xu et al. 2019). However, the expression of CXCL8, an inflammatory cytokine involved in the immune response for urogenital infections, is increased in several cell types in the disease state (**Figure 6E**).

Unexpectedly, the hydrosalpinx sample showed differences in genes associated with cell-to-cell adhesion and ECM-related processes. Drawing from earlier work that quantified molecular changes in chronic kidney disease and fibrosis (Kuppe et al. 2021), we calculated the average expression for each of three sets of genes associated with ECM glycoproteins, collagen, and proteoglycan production, respectively, for each epithelial and stromal subtypes and compared between healthy and disease samples (**Figure 6C**). As expected, the overall expression of the three ECM related gene sets is higher in the stromal cell types than in the epithelial cell subtypes. However, while CE1-4, NSCE2-1 and NCSE2-2 express ECM glycoproteins, collagen, and proteoglycan at low levels, their levels are higher in the disease state (**Figure 6C**). By displaying gene expression heatmaps for the cell type centroids and individual genes, we observed finer patterns of heterogeneity involving cell type-specific changes, and specifically in subsets of genes in the three gene sets (**Figure 6D; Supplemental Table 9**).

Changes in ECM genes in hydrosalpinx raised the possibility that it represents a fibrotic disease. Fibrosis is a known consequence of tissue damage and has been observed in multiple organs. But its underlying molecular and cellular processes in the fallopian tube are unknown (Ajonuma et al. 2005). The TGF-β pathway has been proposed as a main driver of fibrosis in multiple disease processes in many tissues (Biernacka, Dobaczewski, and Frangogiannis 2011). Consistent with the fibrosis model, the hydrosalpinx sample show increased expression of TGF-β pathway effector genes TGFB1, PDGFB, CCN2 (CTGF), and YAP1 in the epithelial and stromal cells (**Figure 6F**). These findings suggest that hydrosalpinx may share a similar mechanism with pathogenic fibrosis, and could potentially benefit from therapies developed for other tissues (Pinto et al. 2017). Unlike previous studies, our scRNA-seq data reveal cell type-specific changes in hydrosalpinx, serving as a resource for identifying more specific biomarkers for earlier intervention and development of targeted therapies.

## DISCUSSION

### A refined cell atlas of the pre-menopausal fallopian tube

We used single-cell RNA-seq to establish a new catalog of cell types, subtypes, and mRNA markers for the healthy human fallopian tube, and identified 12 major cell types (**Figure 7C**). While earlier studies have documented four epithelial cell subtypes: ciliated, secretory, peg, and basal cells, the large cell number and relatively high sequencing depth of our data allowed us to identify 10 subtypes of epithelial cells (**Figure 7B**), thus providing a more refined view of the functional heterogeneity of the pre-menopausal fallopian tube. In particular, the gene expression profiles of these subtypes give a glimpse of multiple progenitor cell populations and their associated transitional or mature cell populations. First, we identified two secretory cell progenitors: (1) an epithelial progenitor (LGR5/PGR) population, NCSE2-5, predicted to give rise to mature secretory cells (i.e., NCSE2-4), and (2) a bipotential progenitor population, NCSE2-2, expressing several EMT markers, predicted to generate ciliated epithelial cells CE1-4 on one hand, and the blood endothelial cells in the stromal compartment on the other (**Figure 7A**). The identification of an EMT expressing progenitor cell population in a simple epithelium is unexpected, since these tissues are believed to be maintained by epithelial stem cells (e.g. LGR5+ cells), which are reliant on active WNT signaling (Clevers 2013). Therefore, in a simple epithelium, the activation of an EMT program is typically only observed in response to wound healing, or during cancer invasion and metastasis in tissue (Nieto et al. 2016). Recently, a study by Hu et. al., analyzed ~1,800 epithelial cells sorted from pre- and post-menopausal fallopian tube samples from ovarian cancer or endometrium cancer patients, and also identified a small number of cells (n~20) bearing an EMT signature (Hu et al. 2020), suggesting that this population can be present in the healthy epithelium, and may be a population vulnerable for cancer initiation. Therefore, the presence of an EMT progenitor pool in the healthy epithelium, reported by us and Hu et. al., raises a question as to how this population is regulated to prevent oncogenic transformation. Curiously, in our dataset we find that the EMT progenitor population (NSCE2-2) expresses PRRX1 (**Figure 3F**), a homeobox factor gene and strong EMT inducer. The loss of PRRX1 is required for tumor initiation and metastasis (Ocaña et al. 2012). Thus, it is possible that in the fallopian tube, PRRX1 serves as a gatekeeper in NSCE2-2 cells, preventing them from adopting a quasi-mesenchymal phenotype, which may lead to tumor initiation, or even subsequent metastasis. Future studies are needed to validate the functional roles of the two progenitor pools in the healthy and disease state by combining genetic reporters with lineage tracing, preferably at the single-cell level.

**Figure 7:**
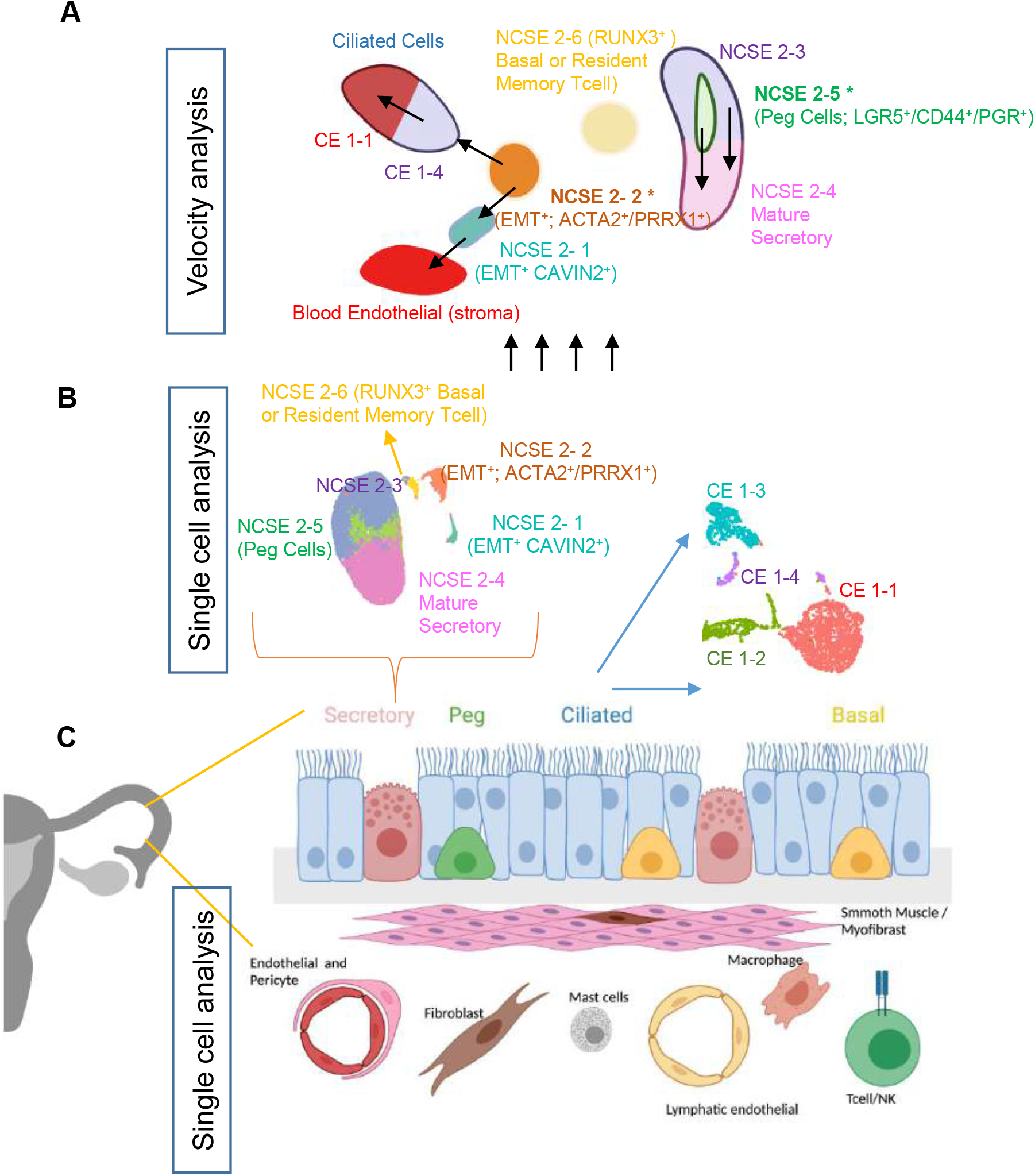
Model for epithelial differentiation trajectory. A. Illustration of the differentiation trajectories of NCSE cells originated from two secretory cell progenitors (NCSE2-2 (ACTA2/PRRX1) and NCSE2-5 (LGR5/PGR)) revealed by velocity analysis. B. ScRNAseq of thousands of fallopian tube cells expands cellular taxonomy of the epithelial cell subtypes from four to ten. C. Schematic of 12 major cell types for healthy human fallopian tubes established by scRNA-seq analysis.

A recent study by Dinh et. al. profiled human fallopian tube samples using 10X Chromium (Dinh et al. 2021). The authors analyzed >50,000 cells, albeit with a shallower sequencing depth than in our dataset (an average of ~850 detected genes per cell, compared to 2,500 in our study), and a wider heterogeneity among samples, which contributed highly varied combinations of major cell types. Importantly, similar to our study, Dinh et al. reported several subtypes among those expressing epithelial and mesenchymal features, which were then designated transitional populations. Pseudotime analysis suggested that one of the transitional populations they observed is a RUNX3^+^ secretory cell type, which was said to have the potential to give rise to ciliated cells. This subtype was not recognizable in our data, for several reasons. First, in our data, RUNX3 is one of the most distinctive markers for the T/NK cells, and is expressed at much lower levels in the other 11 cell types, including the ciliated and non-ciliated epithelial cells (**Figure 1D**). Second, of the four CE subtypes and 6 NCSE subtypes in our data, RUNX3 is expressed highest in NCSE2-6 (**Figure 3G**) (albeit at a much lower level and lower frequency than in the T/NK cells); but NCSE2-6 is not one of the progenitor cells. In our velocity analysis, the RUNX3-expression epithelial cells, NCSE2-6, did not contribute to any other NCSE or CE populations. Genes highly expressed in this population are more consistent with resident memory T-cells (Ardighieri et al. 2014; Hu et al. 2020). In short, our data do not support the existence of a RUNX3-marked progenitor population.

Next, we sought to understand whether hormonal states could influence the progenitor populations we identified. Since high-quality LGR5^+^ antibodies are not available, we could not monitor the protein expression and cell number for NCSE2-5 across the samples covering the cycle. However, we were able to count the number of putative NCSE2-2 cells by an antibody for ACTA2 and compare its frequency across the menstrual cycle for pre-menopausal women, and between pre- and post-menopausal women. The numbers of ACTA2^+^ cells were higher in pre-menopausal women than in post-menopausal women (**Supplemental Figure 3D**), except for one woman on postmenopausal hormone therapy, suggesting that the hormonal milieu may regulate the size of this population. However, the number of ACTA2+ cells that co-express PGR or ESR were very small, suggesting there may be some aspects of the hormonal state that indirectly drives the expansion of this population rather than a direct hormone effect. Future studies should include a larger number of benign samples at varying points in the menstrual cycle to examine this question more carefully.

### Potential role of TGF-β mediated tissue fibrosis in hydrosalpinx

Our analysis of the hydrosalpinx sample revealed changes in both tissue composition (i.e., cell number, **Figure 6B, S6A**) and regulatory states (**Figure 6C–F, S6B-C**). Importantly we identified significant changes in progenitor cell markers between the healthy and disease state. For example, multiple EMT markers, such as THY1, PRRX1, and LGALS, are elevated in the disease state; and we consistently find a higher number of NCSE2-2 cells in hydrosalpinx (**Supplemental Figure 6B, D**). In contrast to EMT markers, LGR5 – an epithelial progenitor marker- levels are significantly reduced in NCSE2-5 cells (**Figure S6B**). However, this is not accompanied by a decrease in NCSE2-5 cell number (**Supplemental Figure 6D**), but rather a rewiring of the transcriptional regulatory network in the disease state.

In addition to changes in progenitor cells, the epithelial and stromal populations express higher levels ECM-related genes (**Figure 6C**), which led us to explore whether the hydrosalpinx disease state may have fibrosis-related tissue damage. Fibrosis has been described for the lung epithelium in idiopathic pulmonary fibrosis (IPF) and many organs in response to tissue damage, whereby epithelial injury activates inflammatory cells, which in turn activate fibroblasts and induce EMT in epithelial cells, leading to the uncontrolled deposition of ECM with abnormal remodeling, and ultimately organ failure (Pinto et al. 2017). In animal models of fibrosis the initial insult is often a brief inflammatory response mediated through the TNF-β pathway, before ultimately progressing to a longer term TGF-β mediated fibrosis (Xu et al. 2019; Palter et al. 2001; Morales et al. 2006; Meng, Nikolic-Paterson, and Lan 2016). In our study we did not observe significant differences in TNF-α related genes, but did detect transcriptomic changes in TGFB1, PDGFB, CCN2, and YAP1 in the hydrosalpinx (**Figure 6E, F**), suggesting that we may have captured a more advanced disease state. To fully test this model would require multiple hydrosalpinx samples at varying time points of disease progression. Nevertheless, several clinical trials are underway for medications targeting individual steps in this fibrosis pathway (Pinto et al. 2017). It is possible that some of these medications could be leveraged as a new pharmacologic therapy for hydrosalpinx and/or tubal factor infertility.

In summary, by sequencing >53K cells from three tubal segments in three healthy subjects we produced a detailed cell atlas of the human fallopian tube, allowing comparisons across anatomic segments of the tube. The identification of 10 subtypes of epithelial cells advanced our knowledge of the maintenance and dynamic regulation of the FT epithelium and established a baseline to study changes of cell number and cell state across many types of biological variation: among tubal segments, during the hormone cycle, from pre-menopausal to peri- and post-menopausal, and for diseases such as tubal factor infertility and ovarian cancer. Here we analyzed hydrosalpinx as one example of such comparisons; and the same approach can be extended to other variations of interest. Furthermore, by using computational trajectory analysis we were able to extract information regarding the direction of cell differentiation, and this allowed us to define multiple putative progenitor cells, which will likely be a focus of future studies of tissue homeostatic as well as its dysregulation in diseases.

## Acknowledgements

We thank members of the Hammoud, and Li Labs for scientific discussions and manuscript comments. This research was supported by National Institute of Health (NIH) training grants T32-HD079342 (A.J.), T32-GM70449 (D.F.H.), fellowship F31-HD106626 (A.J.) and Chan Zuckerberg Foundation Grant CZF2019-002428 (A.S., E.E.M., J.Z.L. and S.S.H.).

## Supplemental Figure Legends

**Figure S1. Assessing reproducibility of discovered cell types in healthy human fallopian tubes across tubal segments and across subjects.**

A. Consistent clustering patterns across the 3 segments from FT1. Shown are UMAP projections of the 1,861 fimbria cells, 4,944 ampulla cells, and 3,722 isthmus cells, colored by the 11 clusters independently obtained for the 3 segments.

B. Heatmap of rank correlation coefficients for pairs of cluster centroids across 11 clusters for each of the 3 segments, showing that most cell types were consistently observed across segments.

C. After combining the three segments, rank correlation coefficients of cluster centroid pairs showed consistency across fallopian tube samples from the three healthy subjects. Clustering was performed separately for each subject with cells combined across segments.

D. Visualization of the cells from each of the 3 samples in global UMAP projection, with one sample colored and the other two in grey.

E. Cell type composition of the 12 major cell types, compared across the 3 fallopian tube segments and the 3 healthy subjects.

**Figure S2: Ciliated cell subtypes.**

A. Consistent representation of ciliated cell subtypes in the 3 healthy fallopian tube samples. Shown are the same UMAP projections, with each panel displaying cells from one subject in color, on the grey background of cells from the other two subjects.

B. Comparison of “cell size factor”, the per-cell count of detected transcripts, over the 4 subtypes.

C. Comparison of the percentage of mitochondria encoded RNA in all detected RNA for the cells.

D. Heatmap of rank correlation coefficients between individual cells (in rows) and five cluster centroids (in columns). From top to bottom are 1,452 ciliated cells of fimbria2 (ordered by their assignments into 4 subtypes) and 1,424 soup-like cells, with <100 UMIs per cell. From left to right are the 4 CE subtype centroids and the soup centroid (see Methods). Correlation values were calculated by using 2000 highly variable genes among the 4 CE centroids.

E. Protein expression of CDKN2A in a fallopian tube section. Arrow: CDKN2A^+^ cells. Courtesy of Human Protein Atlas. https://www.proteinatlas.org/ENSG00000147889-CDKN2A/tissue/fallopian+tube#img

**Figure S3: Secretory cell subtypes.**

A. Reproducibility of epithelial cell subtypes in the 3 healthy fallopian tube samples. Shown are the same UMAP projections, with each panel displaying cells from one subject in color, on the grey background of cells from the other two subjects.

B. Cell number and percentage of the 6 NCSE subtypes, for each of the 3 fallopian tube samples from subjects healthy.

C. Joint distribution of expression levels for 4 pairs of genes over individual cells, colored by the 6 NCSE subtypes.

D. Comparison of the percentage of CE cells (top), NCSE 2-2 cells (middle), and NCSE 2-6 cells (bottom), in the 3 segments of the fallopian tube, for 11 subjects (left to right), covering different menopausal stages. The diseased sample, FT3, is indicated by the box. CE, NCSE2-2 and NCSE2-6 cells were defined using a combination of EPCAM^+^/CAPS^+^/FOXJ1^+^, EPCAM^+^/ACTA2^+^/FOXJ1^-^, EPCAM^+^/RUNX3^+^/ CD44^+^, respectively. Percent values were calculated per 1000 cells counted randomly in four quadrants of the tissue cross-section.

E. IF co-staining of ACTA2 in the fallopian tube epithelium with estrogen receptor (top) or progesterone receptor (bottom).

F. Expression levels of genes encoding hormone receptors in the 6 NCSE subtypes.

**Figure S4: Stromal cells subset analysis.**

A. Consistent representation of the six stromal cell subtypes in the 3 healthy fallopian tube samples. Shown are the same UMAP projections, with each panel displaying cells from one subject in color, on the grey background of cells from the other two subjects.

B. IF staining in the fallopian tube epithelium using the antibodies against a specific marker for each of the 6 stromal cell types. PRXX1, NCAM, and VWF are courtesy of Human Protein Atlas. PRXX1:. MCAM: https://www.proteinatlas.org/ENSG00000076706-MCAM/tissue/fallopian+tube#img. VWR: https://www.proteinatlas.org/ENSG00000110799-VWF/tissue/fallopian+tube#img.

**Figure S5: Gene expression values vary across segments.**

A. Expression levels of select markers differentially expressed across the 3 tubal segments, for ciliated cells (top) and non-ciliated secretory epithelial cells (bottom).

B. Expression of ALDH1A1 and ALDH1A2 across 3 tubal segments, compared across the 4 ciliated and 6 non-ciliated secretory epithelial cell subtypes.

**Figure S6: Comparison between a hydrosalpinx sample and healthy fallopian tube samples.**

A. Comparison of the percentage of MKI67^+^ cells between healthy and diseased samples, for 4 CE subtypes, 6 NCSE subtypes and 6 stromal cell types.

B. Expression levels of EMT (left) and stemness (right) markers between healthy (orange) and disease (blue) samples, for CE cells, NCSE subtypes and (for ALDH1A1) the stromal cell types.

C. Expression levels of ovarian cancer-related genes compared between healthy (orange) and disease (blue) samples, for the CE cells and NCSE subtypes.

D. Cell number and percentage of the 4 CE subtypes and 6 NCSE subtypes, compared between the 3 healthy subjects and the diseased subject.

## Notes

### Competing Interest Statement

The authors have declared no competing interest.

