## Supplemental Figures for "Cellular heterogeneity of human fallopian tubes in normal and hydrosalpinx disease states identified by scRNA-seq"

#### Supplemental Figure 1

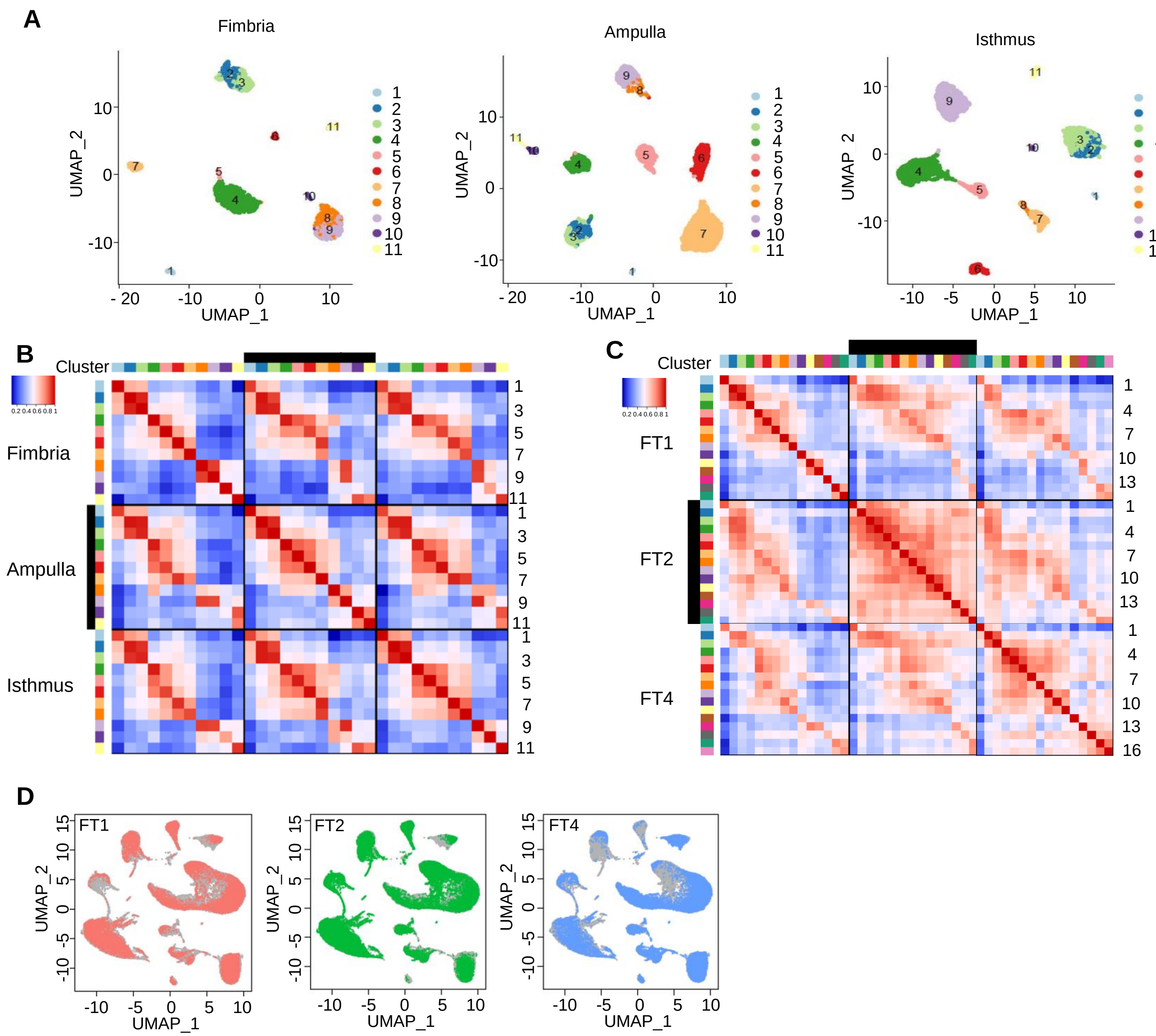


**E**

| | | 1 | 2 | 3 | 4 | 5 | 6 | 7 | 8 | 9 | 10 | 11 | 12 | Total |
|---|---|---|---|---|---|---|---|---|---|---|---|---|---|---|
| FT1 | F | 2.3% | 22.7% | 8.9% | 24.1% | 0.6% | 0.9% | 2.4% | 5.3% | 0.6% | 27.4% | 1.3% | 3.1% | 1861 |
| | A | 0.7% | 13.5% | 8.6% | 33.8% | 10.5% | 7.1% | 8.9% | 1.1% | 0.7% | 11.2% | 0.2% | 2.6% | 4944 |
| | I | 0.9% | 27.5% | 6.6% | 18.8% | 5.6% | 4.2% | 7.1% | 1.6% | 0.8% | 22.7% | 1.2% | 2.5% | 3722 |
| FT2 | F | 15.6% | 40.9% | 4.7% | 20.4% | 0.4% | 0.3% | 2.5% | 2.3% | 0.0% | 6.7% | 3.5% | 2.6% | 9332 |
| | A | 4.4% | 48.3% | 3.0% | 2.5% | 12.7% | 8.3% | 11.6% | 0.6% | 0.0% | 2.4% | 3.4% | 2.7% | 8021 |
| | I | 1.1% | 28.3% | 6.0% | 4.7% | 19.6% | 9.7% | 20.0% | 0.7% | 0.1% | 2.8% | 4.0% | 3.1% | 8043 |
| FT4 | F | 4.2% | 16.2% | 18.2% | 21.3% | 0.2% | 0.0% | 0.3% | 13.5% | 0.2% | 14.7% | 3.8% | 6.9% | 8200 |
| | A | 3.0% | 9.2% | 22.6% | 19.3% | 2.6% | 0.6% | 2.3% | 11.2% | 1.1% | 13.2% | 2.2% | 12.8% | 2181 |
| | I | 1.4% | 5.5% | 19.0% | 25.2% | 2.9% | 0.5% | 2.7% | 6.6% | 5.0% | 18.8% | 5.0% | 6.7% | 7072 |
| | Total | 2.3% | 22.7% | 8.9% | 24.1% | 0.6% | 0.9% | 2.4% | 5.3% | 0.6% | 27.4% | 1.3% | 3.1% | 53376 |

**Supplemental Figure 1: Assessing reproducibility of discovered cell types in healthy human fallopian tubes across tubal segments and across subjects.**

A. Consistent clustering patterns across the 3 segments from FT1. Shown are UMAP projections of the 1,861 fimbria cells, 4,944 ampulla cells, and 3,722 isthmus cells, colored by the 11 clusters independently obtained for the 3 segments.

B. Heatmap of rank correlation coefficients for pairs of cluster centroids across 11 clusters for each of the 3 segments, showing that most cell types were consistently observed across segments.

C. After combining the three segments, rank correlation coefficients of cluster centroid pairs showed consistency across fallopian tube samples from the three healthy subjects. Clustering was performed separately for each subject with cells combined across segments.

D. Visualization of the cells from each of the 3 samples in global UMAP projection, with one sample colored and the other two in grey.

E. Cell type composition of the 12 major cell types, compared across the 3 fallopian tube segments and the 3 healthy subjects.

### Supplemental Figure 2

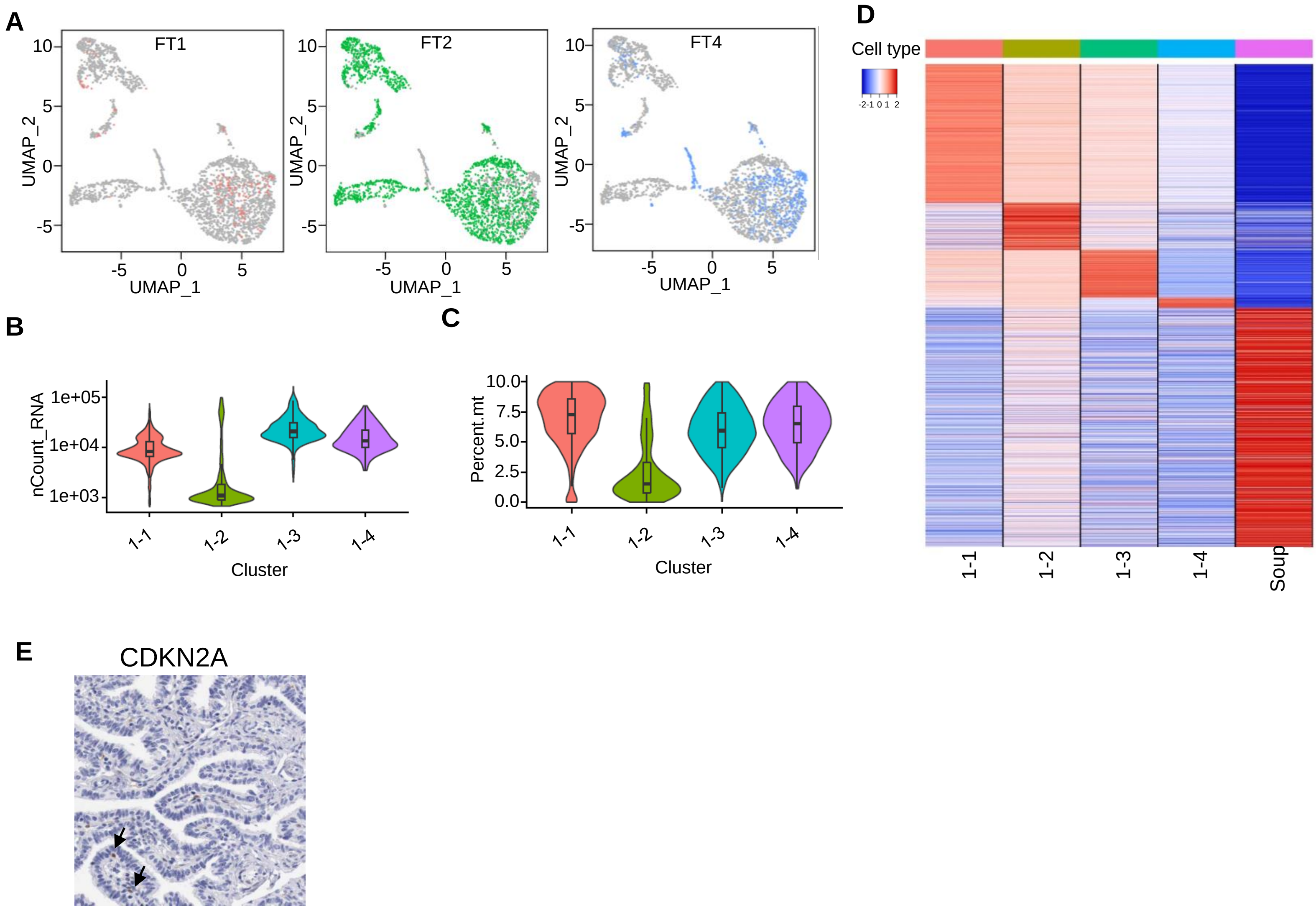

**Supplemental Figure 2: Ciliated cell subtypes.**

A. Consistent representation of ciliated cell subtypes in the 3 healthy fallopian tube samples. Shown are the same UMAP projections, with each panel displaying cells from one subject in color, on the grey background of cells from the other two subjects.

B. Comparison of "cell size factor", the per-cell count of detected transcripts, over the 4 subtypes.

C. Comparison of the percentage of mitochondria encoded RNA in all detected RNA for the cells.

D. Heatmap of rank correlation coefficients between individual cells (in rows) and five cluster centroids (in columns). From top to bottom are 1,452 ciliated cells of fimbria2 (ordered by their assignments into 4 subtypes) and 1,424 soup-like cells, with <100 UMIs per cell. From left to right are the 4 CE subtype centroids and the soup centroid (see Methods). Correlation values were calculated by using 2000 highly variable genes among the 4 CE centroids.

E. Protein expression of CDKN2A in a fallopian tube section. Arrow: $CDKN2A^{+}$ cells. Courtesy of Human Protein Atlas. https://www.proteinatlas.org/ENSG00000147889-CDKN2A/tissue/fallopian+tube#img

### Supplemental Figure 3

A

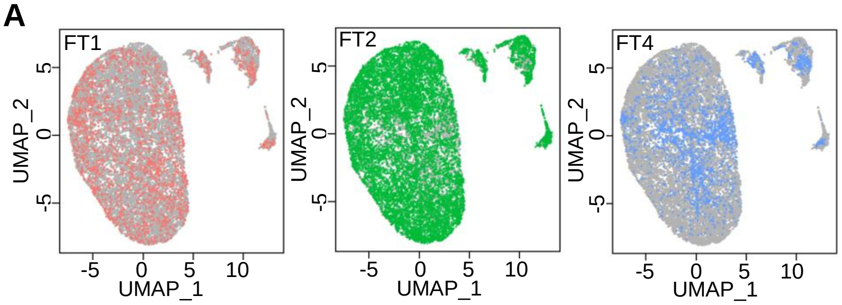


C

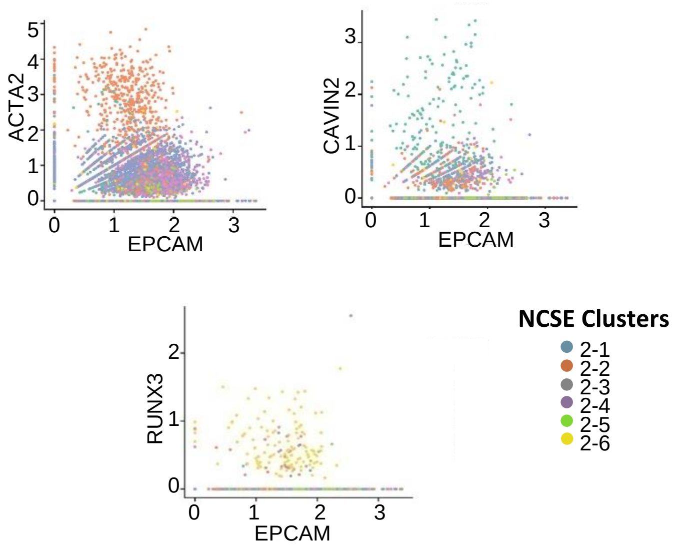


B

Number of Cells (%)

| Cluster | 1 | 2 | 3 | 4 | 5 | 6 | Total |
|---|---|---|---|---|---|---|---|
| FT1 | 34 (1.6) | 121 (5.7) | 935 (44.2) | 932 (44.1) | 44 (2.1) | 47 (2.2) | 2113 |
| FT2 | 200 (2.0) | 605 (6.1) | 5011 (50.2) | 3942 (39.5) | 21 (0.2) | 194 (1.9) | 9973 |
| FT4 | 34 (1.8) | 142 (7.4) | 576 (30.1) | 605 (31.6) | 484 (25.3) | 75 (3.9) | 1916 |

D

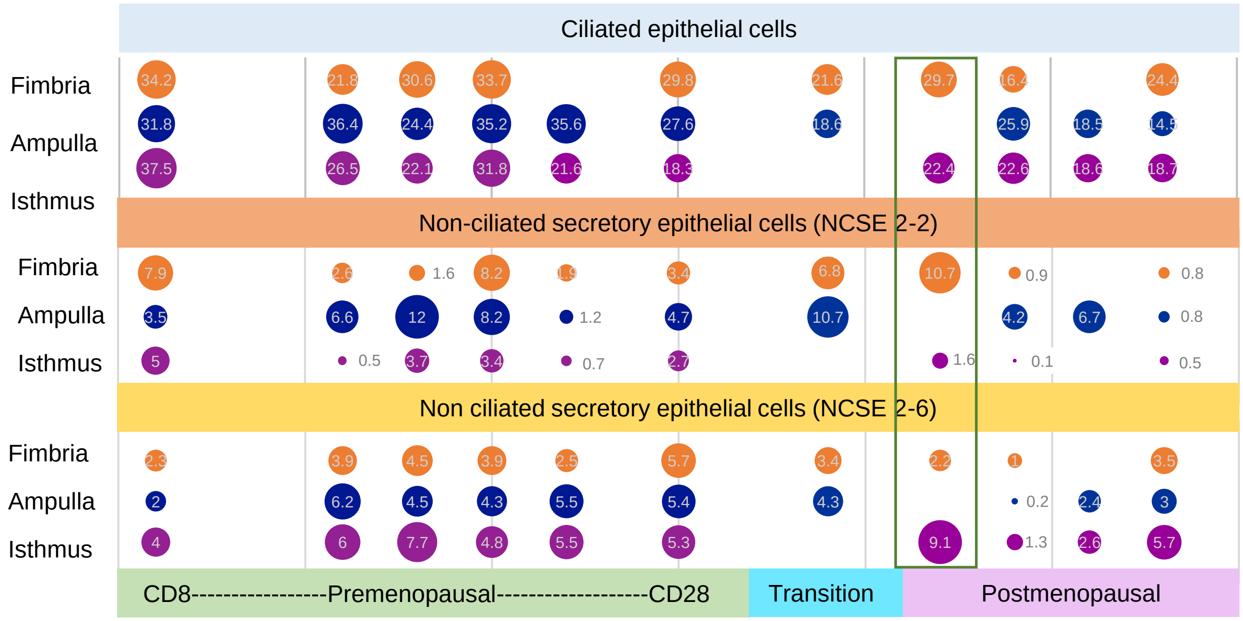


E

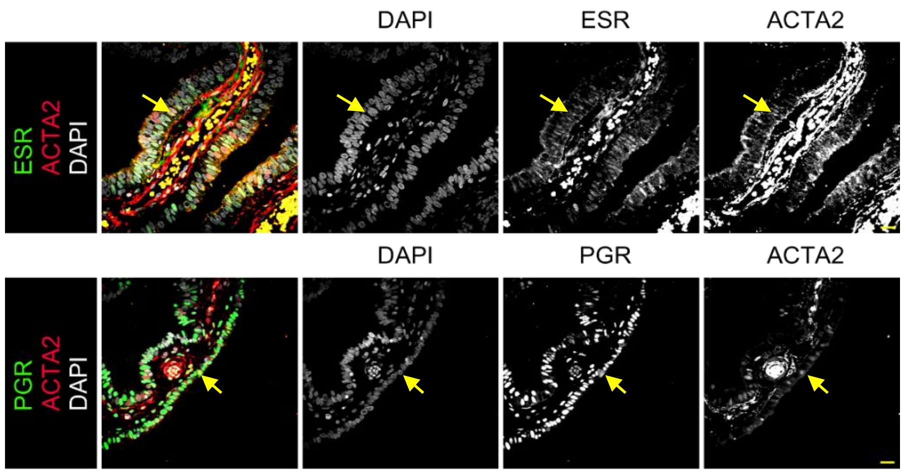


F

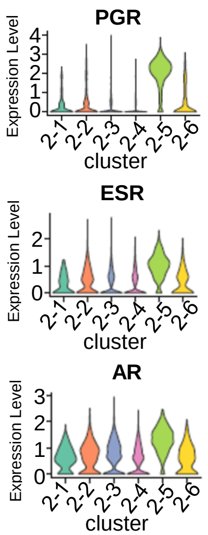

**Supplemental Figure 3: Secretory cell subtypes.**

A. Reproducibility of epithelial cell subtypes in the 3 healthy fallopian tube samples. Shown are the same UMAP projections, with each panel displaying cells from one subject in color, on the grey background of cells from the other two subjects.

B. Cell number and percentage of the 6 NCSE subtypes, for each of the 3 fallopian tube samples from subjects healthy.

C. Joint distribution of expression levels for 4 pairs of genes over individual cells, colored by the 6 NCSE subtypes.

D. Comparison of the percentage of CE cells (top), NCSE 2-2 cells (middle), and NCSE 2-6 cells (bottom), in the 3 segments of the fallopian tube, for 11 subjects (left to right), covering different menopausal stages. The diseased sample, FT3, is indicated by the box. CE, NCSE2-2 and NCSE2-6 cells were defined using a combination of $EPCAM^{+}$/$CAPS^{+}$/$FOXJ1^{+}$, $EPCAM^{+}$/$ACTA2^{+}$/$FOXJ1^{-}$, $EPCAM^{+}$ /$RUNX3^{+}$/ $CD44^{+}$, respectively. Percent values were calculated per 1000 cells counted randomly in four quadrants of the tissue cross-section.

E. IF co-staining of ACTA2 in the fallopian tube epithelium with estrogen receptor (top) or progesterone receptor (bottom).

F. Expression levels of genes encoding hormone receptors in the 6 NCSE subtypes.

### Supplemental Figure 4

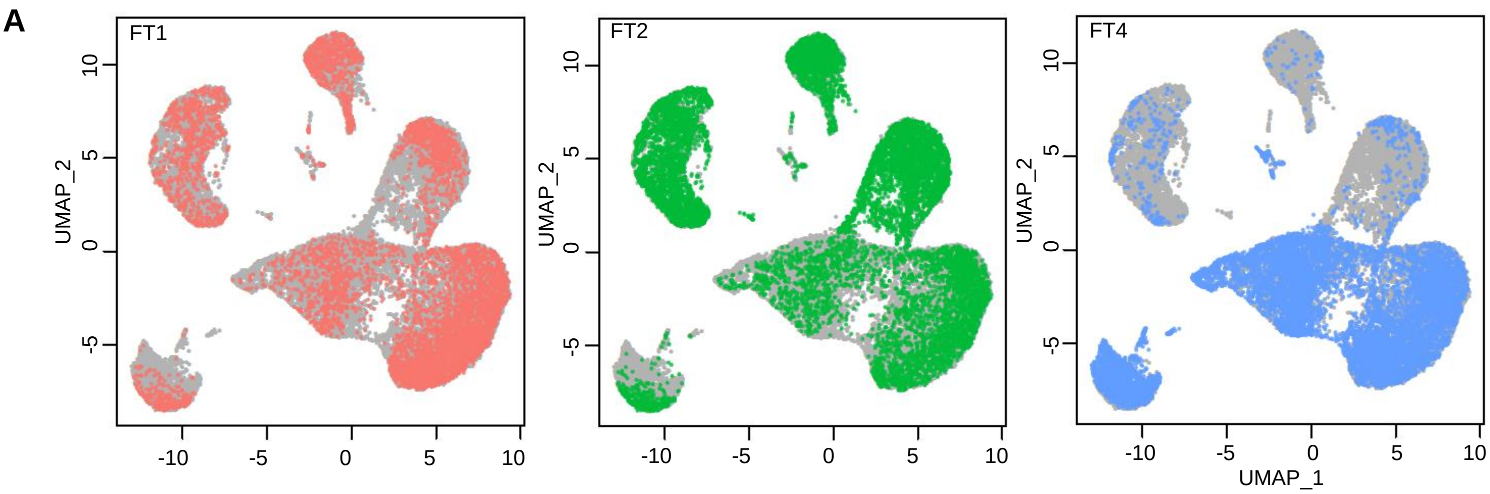


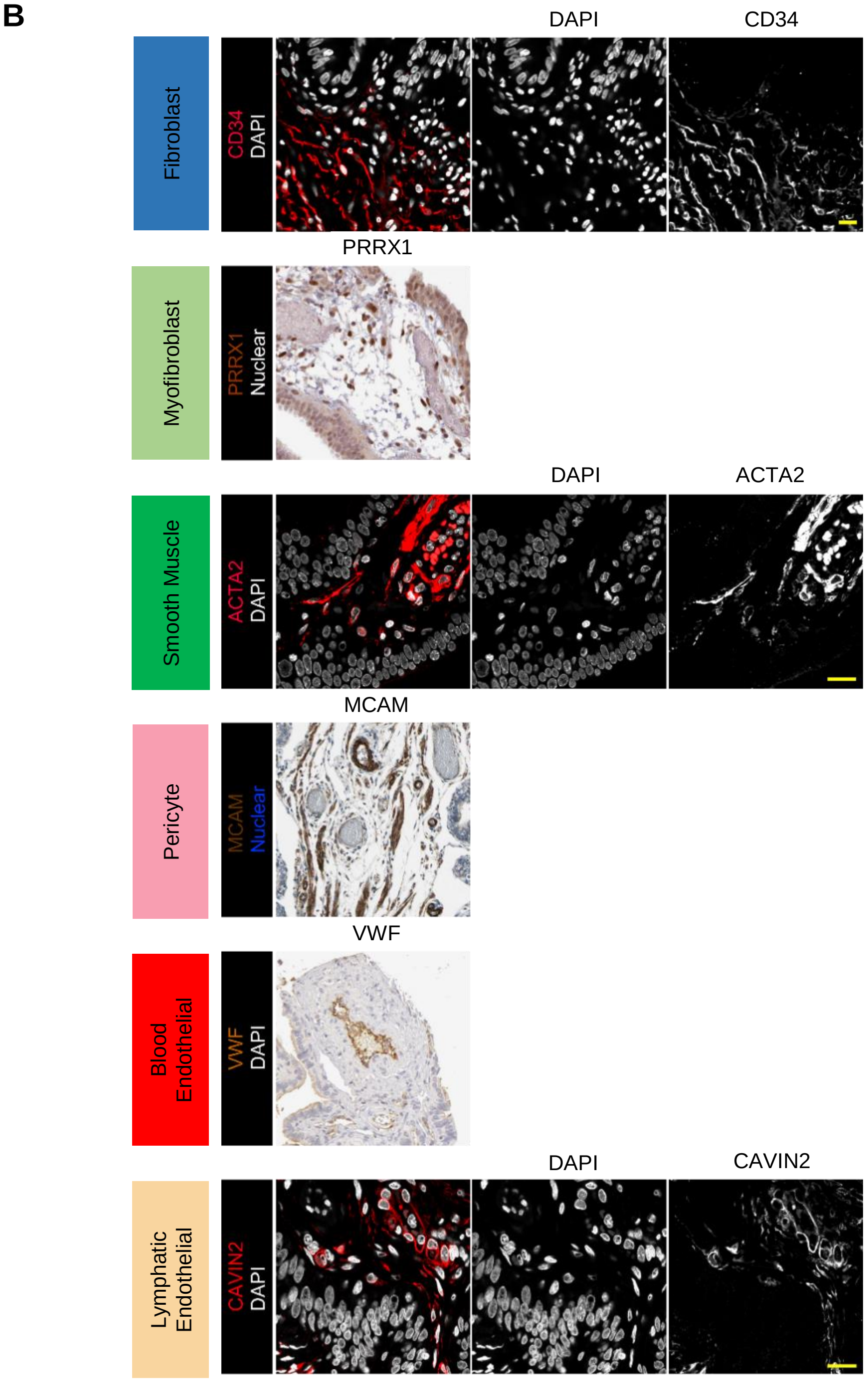

**Supplemental Figure 4: Stromal cells subset analysis.**

A. Consistent representation of the six stromal cell subtypes in the 3 healthy fallopian tube samples. Shown are the same UMAP projections, with each panel displaying cells from one subject in color, on the grey background of cells from the other two subjects.

B. IF staining in the fallopian tube epithelium using the antibodies against a specific marker for each of the 6 stromal cell types. PRXX1, NCAM, and VWF are courtesy of Human Protein Atlas.
PRXX1: https://www.proteinatlas.org/ENSG00000116132-PRRX1/tissue/fallopian+tube#img.
MCAM: https://www.proteinatlas.org/ENSG00000076706-MCAM/tissue/fallopian+tube#img.
VWR: https://www.proteinatlas.org/ENSG00000110799-VWF/tissue/fallopian+tube#img.

### Supplemental Figure 5

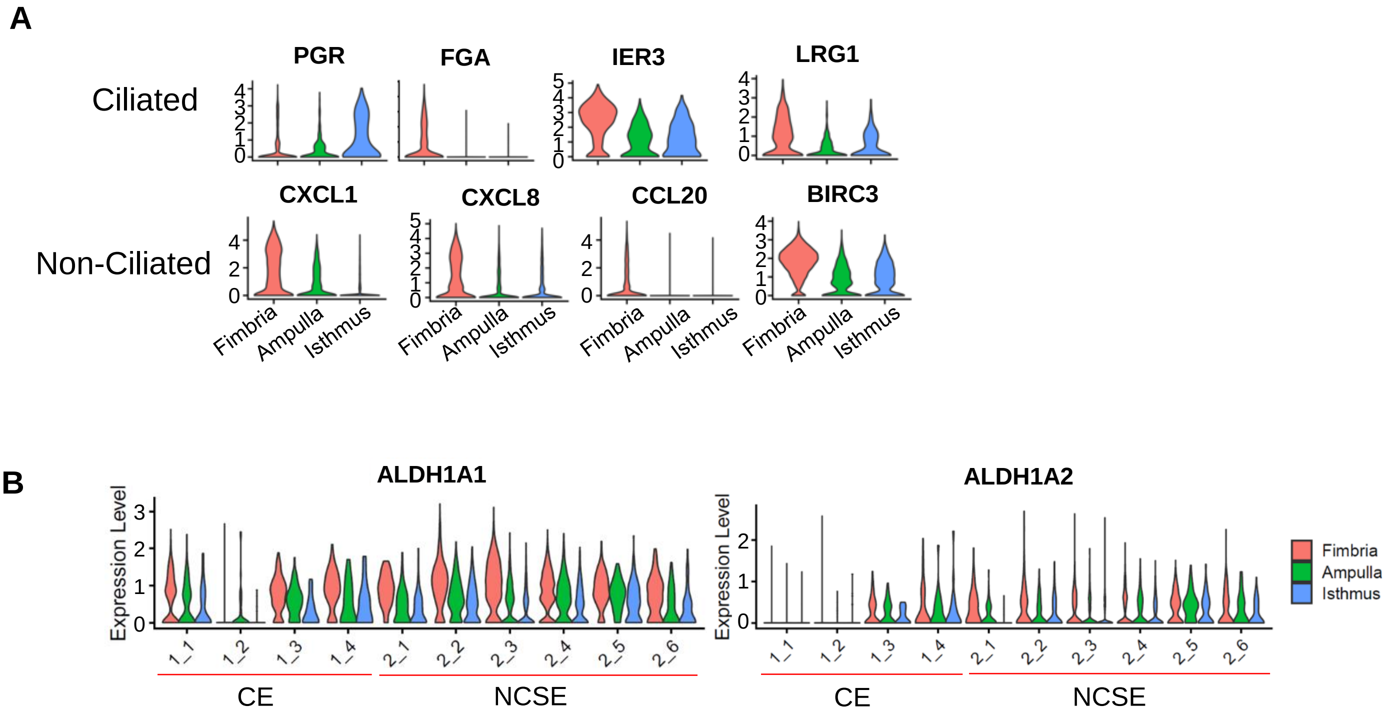

**Supplemental Figure 5: Gene expression values vary across segments.**

A. Expression levels of select markers differentially expressed across the 3 tubal segments, for ciliated cells (top) and non-ciliated secretory epithelial cells (bottom).

B. Expression of ALDH1A1 and ALDH1A2 across 3 tubal segments, compared across the 4 ciliated and 6 non-ciliated secretory epithelial cell subtypes.

#### Supplemental Figure 6

**A**

Fraction of MKI67 positive cells

| | | Healthy (%) | Disease (%) |
|---|---|---|---|
| Ciliated Cells | 1_1 | 0.07 | 0.23 |
| | 1_2 | 2.58 | 0.00 |
| | 1_3 | 0.58 | 1.92 |
| | 1_4 | 0.00 | 1.59 |
| NonCiliated Cells | 2_1 | 0.00 | 7.14 |
| | 2_2 | 0.81 | 5.33 |
| | 2_3 | 0.49 | 6.31 |
| | 2_4 | 0.22 | 3.57 |
| | 2_5 | 1.82 | 4.63 |
| | 2_6 | 1.58 | 2.45 |
| | Fib | 0.24 | 0.11 |
| | MyoFib | 0.01 | 0.11 |
| | SM | 0.00 | 0.23 |
| | Pericyte | 0.05 | 0.00 |
| | B- Endo | 0.03 | 1.26 |
| | L- Endo | 0.26 | 0.22 |

**B**

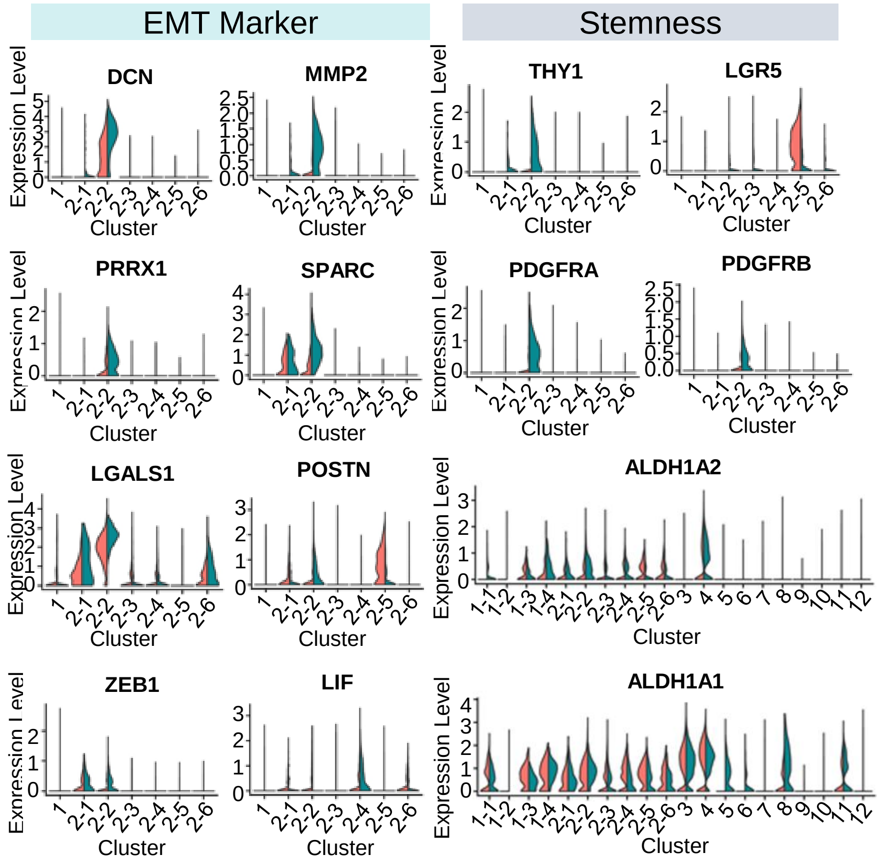


**C**

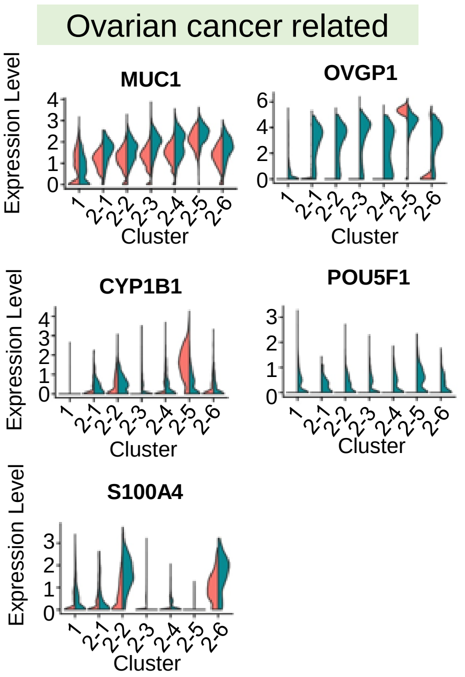


**D**

| | 1_1 | 1_2 | 1_3 | 1_4 | 2_1 | 2_2 | 2_3 | 2_4 | 2_5 | 2_6 | Total |
|---|---|---|---|---|---|---|---|---|---|---|---|
| Number of cells | | | | | | | | | | | |
| Disease | 444 | 0 | 52 | 63 | 70 | 300 | 1299 | 224 | 410 | 204 | 3066 |
| Healthy | 1466 | 387 | 513 | 144 | 268 | 868 | 6522 | 5479 | 549 | 316 | 16512 |
| Total | 1910 | 387 | 565 | 207 | 338 | 1168 | 7821 | 5703 | 959 | 520 | 19578 |
| Constitutes | | | | | | | | | | | |
| Disease | 14.5% | 0.0% | 1.7% | 2.1% | 2.3% | 9.8% | 42.4% | 7.3% | 13.4% | 6.7% | 1 |
| Healthy | 8.9% | 2.3% | 3.1% | 0.9% | 1.6% | 5.3% | 39.5% | 33.2% | 3.3% | 1.9% | 1 |

**Supplemental Figure 6: Comparison between a hydrosalpinx sample and healthy fallopian tube samples.**

A. Comparison of the percentage of $MKI67^{+}$ cells between healthy and diseased samples, for 4 CE subtypes, 6 NCSE subtypes and 6 stromal cell types.

B. Expression levels of EMT (left) and stemness (right) markers between healthy (orange) and disease (blue) samples, for CE cells, NCSE subtypes and (for ALDH1A1) the stromal cell types.

C. Expression levels of ovarian cancer-related genes compared between healthy (orange) and disease (blue) samples, for the CE cells and NCSE subtypes.

D. Cell number and percentage of the 4 CE subtypes and 6 NCSE subtypes, compared between the 3 healthy subjects and the diseased subject.
